# Conservation of dichromatin organization along regional centromeres

**DOI:** 10.1101/2023.04.20.537689

**Authors:** Danilo Dubocanin, Gabrielle A. Hartley, Adriana E. Sedeno Cortes, Yizi Mao, Sabrine Hedouin, Jane Ranchalis, Aman Agarwal, Glennis A. Logsdon, Katherine M. Munson, Taylor Real, Benjamin J. Mallory, Evan E. Eichler, Sue Biggins, Rachel J. O’Neill, Andrew B. Stergachis

## Abstract

The focal attachment of the kinetochore to the centromere is essential for genome maintenance, yet the highly repetitive nature of satellite regional centromeres, such as those in humans, limits our understanding of their chromatin organization. We demonstrate that single-molecule chromatin fiber sequencing (Fiber-seq) can uniquely co-resolve kinetochore and surrounding chromatin architectures along point centromeres, revealing largely homogeneous single-molecule kinetochore occupancy along each chromosome. In contrast, extension of Fiber-seq to regional satellite centromeres exposed marked per-molecule heterogeneity in their chromatin organization. Regional CENP-A-marked centromere cores uniquely contain a dichotomous chromatin organization (dichromatin) composed of compacted nucleosome arrays punctuated with highly accessible chromatin patches. CENP-B occupancy phases dichromatin to the underlying alpha-satellite repeat within centromere cores, but is not necessary for dichromatin formation. Centromere core dichromatin is a conserved feature between humans despite the marked divergence of their underlying alpha-satellite organization and is similarly a conserved feature along regional centromeres that lack satellite repeats in gibbon. Overall, the chromatin organization of regional centromeres is defined by marked per-molecule heterogeneity, likely buffering kinetochore attachment against sequence and structural variability within regional centromeres.

**Highlights:**

- Dichotomous accessible and compacted chromatin (dichromatin) marks centromere cores
- Highly accessible chromatin patches punctuate sites of kinetochore attachment
- Dichromatin can form irrespective of CENP-B occupancy
- Conservation within centromeres is mediated at the level of chromatin, not DNA

## Introduction

Centromeres can range from ‘point’ centromeres, which contain a single well positioned CENP-A nucleosome, to ‘regional’ centromeres, which include numerous CENP-A nucleosomes often embedded within tandemly repeated sequences. For example, *Saccharomyces cerevisiae* centromeres are comprised of ∼125 bp point centromeres built on a single CENP-A nucleosome, whereas human centromeres are comprised of regional centromeres that occupy ∼171 bp alpha-satellite repeat units organized into higher-order repeats (HORs) that can span several megabases on each chromosome. These alpha-satellite HORs serve as the genetic substrate for kinetochore attachment to the human genome within the centromere core^1,2^, and kinetochore interactions within the centromere core are modulated by several DNA-binding proteins, including the sequence-specific DNA binding protein CENP-B^3,4^ and the histone H3 variant CENP-A^5^. Imaging and genomic studies have demonstrated that kinetochore attachment is limited to only a small portion of the alpha-satellite HOR array, which is marked by CENP-A and CENP-B occupancy along the autosomes and X chromosome^6,7^ and hypo-CpG methylation^8–10^ (*i.e.,* the centromere core or ‘centromere dip region [CDR]’). However, *in vivo* chromatin compaction and organization within this region remains largely unresolved, with somewhat contradictory chromatin features appearing to localize to the centromere core. For example, the centromere core contains a unique patterning of histones referred to as ‘centro-chromatin’, which is comprised of both centromere-specific CENP-A nucleosomes, as well as H3 nucleosomes dimethylated on Lys4 (H3K4me2)^11^ that are often associated with euchromatic portions of the genome^12^. Despite the presence of these H3K4me2 histone, the centromere core is thought to contain condensed chromatin arrays *in vivo*^13^, which are proposed to arise from CENP-A mediated nucleosome condensation^14^ as well as other kinetochore proteins bridging and compacting neighboring nucleosomes^15–17^.

Resolving exactly how chromatin is organized along individual chromatin fibers within the centromere as well as the relationship between kinetochore binding and chromatin organization has been challenging due to the highly repetitive DNA content of alpha-satellite HORs^13^. Specifically, centromeres are systematically unresolved in most reference genomes, and short-read sequencing-based chromatin profiling methods cannot uniquely resolve chromatin architectures across these repetitive DNA arrays (**Figure S1A**). In addition, short-read sequencing-based chromatin profiling methods are inherently unable to resolve how multiple chromatin features are positioned along a single chromatin fiber, as these methods inherently massively fragment chromatin fibers in the process of studying them.

The completion of the first telomere-to-telomere (T2T) human reference genome^18^ in combination with emerging methyltransferase-based single-molecule long-read chromatin profiling methods^19–22^ opens the possibility of studying chromatin features across centromeres at single-molecule and single-nucleotide resolution. Specifically, non-specific adenine methyltransferase (m6A-MTase)-based chromatin profiling methods (i.e., Fiber-seq) enable nucleotide-precise mappings of chromatin accessibility, protein occupancy, nucleosome positioning, and CpG methylation along multi-kilobase chromatin fibers^19,23^ (**Figure S1B and S1C**). Fiber-seq utilizes a non-specific m6A-MTase^24^ to stencil the chromatin architecture of individual multi-kilobase fibers onto their underlying DNA templates via methylated adenines (**Figure 1A**), which is a non-endogenous DNA modification in humans^25,26^. The genetic and chromatin architecture of each fiber is directly read using highly accurate single-molecule PacBio HiFi long-read DNA sequencing, which is capable of accurately distinguishing m6A-and mCpG-modified bases at single-nucleotide resolution^23,27^.

**Figure 1.**
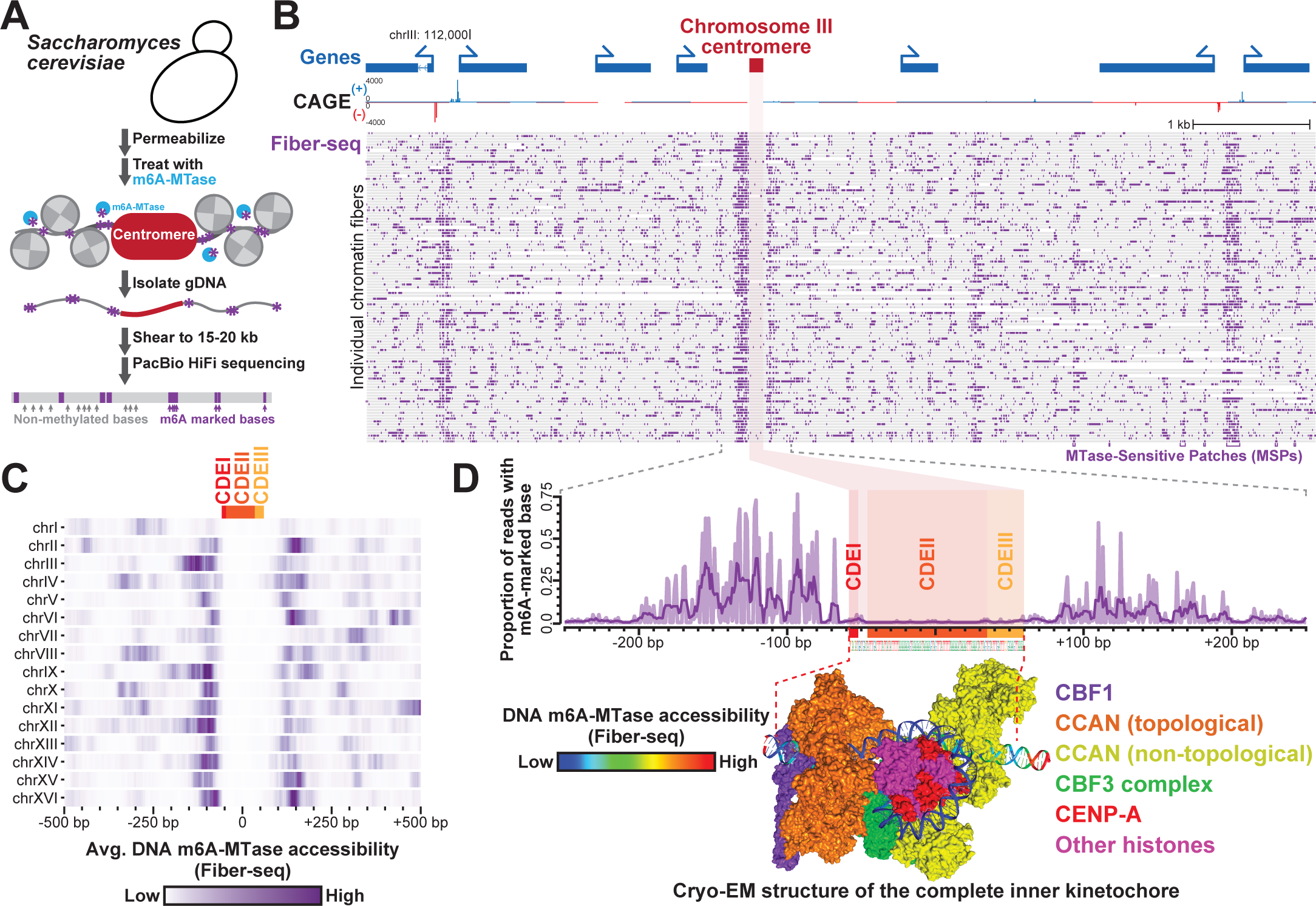
Resolution of kinetochore and surrounding chromatin architectures along point centromeres. (A) Schematic for mapping the chromatin occupancy of point centromeres within the yeast *S. cerevisiae* using Fiber-seq. (B) Locus displaying single-molecule chromatin architectures of the *S. cerevisiae* chromosome III point centromere, alongside CAGE data and gene annotations. Note the large patches of chromatin accessibility at the CAGE-positive gene promoters, as well as immediately adjacent to the centromere. (C) Heatmap displaying average Fiber-seq m6A signal surrounding each of the *S. cerevisiae* point centromeres. (D) (top) Zoom-in of the per-nucleotide Fiber-seq signal of the *S. cerevisiae* chromosome III point centromere in relation to the CDEI, CDEII and CDEIII elements. (bottom) Cryo-EM structure of the yeast complete *S. cerevisiae* inner kinetochore bound to the chromosome III point centromere (PDB 8OW0) with the DNA colored according to the m6A-MTase sensitivity measured via Fiber-seq on *S. cerevisiae*.

## Results

### Single-molecule chromatin architectures of kinetochore-bound point centromeres

To map the chromatin architecture of centromeres at single-molecule and near single-nucleotide resolution, we first applied Fiber-seq to budding yeast *Saccharomyces cerevisiae* (**Fig. 1A**), which contain a point centromere along each chromosome^28,29^. Similar to humans, the yeast inner kinetochore directly interacts with DNA via a centromere-associated inner kinetochore (CCAN) complex, which contains the histone H3 variant CENP-A (Cse4)^28^ in addition to the yeast-specific centromere binding factor 1 (Cbf1)^30^ and the yeast-specific multiprotein complex CBF3^31^. Each yeast centromere is organized into three centromere DNA elements (CDEs) (CDEI, CDEII, and CDEIII), and cryo-EM structures and nuclease digestions of the yeast CCAN complex have shown that the yeast CCAN embeds ∼160 bp of DNA within this complex^32,33^. Application of Fiber-seq to asynchronously growing *Saccharomyces cerevisiae* cells revealed a discrete change in chromatin structure immediately overlapping the CDEI-III elements (**Fig. 1B, 1C**). Specifically, >90% of chromatin fibers overlapping the yeast centromere contained a well-positioned MTase-protected region overlapping the CDEI-III elements. This MTase-protected region was significantly larger than the standard nucleosome footprint size observed genome-wide, perfectly aligned with the cryo-EM structure of the yeast inner kinetochore (**Fig. 1D**), and was disrupted upon Cse4 destabilization^34^. Notably, yeast centromeres were frequently flanked by MTase sensitive patches (MSP) of chromatin that mirror the size of traditional gene regulatory elements (**Fig. 1B, 1C**), indicating that the yeast CCAN is associated with marked alterations in the local chromatin architecture. Together, these findings reveal that Fiber-seq can resolve the chromatin architecture of the CCAN at single-molecule and near single-nucleotide resolution and that individual fibers demonstrate homogeneous CCAN occupancy that frequently abuts patches of accessible chromatin.

### Dichotomous chromatin marks the centromere core in humans

We next sought to investigate the chromatin architecture of human centromeres by applying Fiber-seq to the human hydatidiform mole CHM13 cell line^35^, the same line used for constructing the first human telomere-to-telomere (T2T) reference genome^9,18^. We observed that coupling long-read epigenome maps with paired fully sequenced and assembled genome maps from the same individual enabled us to uniquely map the chromatin architecture of all centromeres in CHM13 cells (**Figure 2**), overcoming the substantial diversity between individuals^36^ that is enriched within these highly repetitive genomic regions. Notably, in stark contrast to the yeast point centromeres, CHM13 regional centromeres lacked homogeneously positioned chromatin (**Fig. 2A**). However, similar to yeast, CHM13 centromere cores contained markedly altered chromatin structures not observed elsewhere in the human genome. Specifically, the nucleosome architecture of CHM13 centromere cores diverged from that observed elsewhere in the human genome, containing mononucleosome footprints markedly smaller than the average nucleosome footprint size outside of the centromere core (**Figure S1D**), consistent with a large population of these centromere core mononucleosomes containing CENP-A, which is known to wrap only 121 bp of DNA in humans^13,37,38^. In addition, the centromere core contained some of the most compacted nucleosome arrays within the human genome, with nearly 50% of all nucleosome footprints within the centromere core being di-nucleosomal in size (i.e. >210 bp), consistent with prior observations^39^ (**Fig. 2B and 2C**).

**Figure 2.**
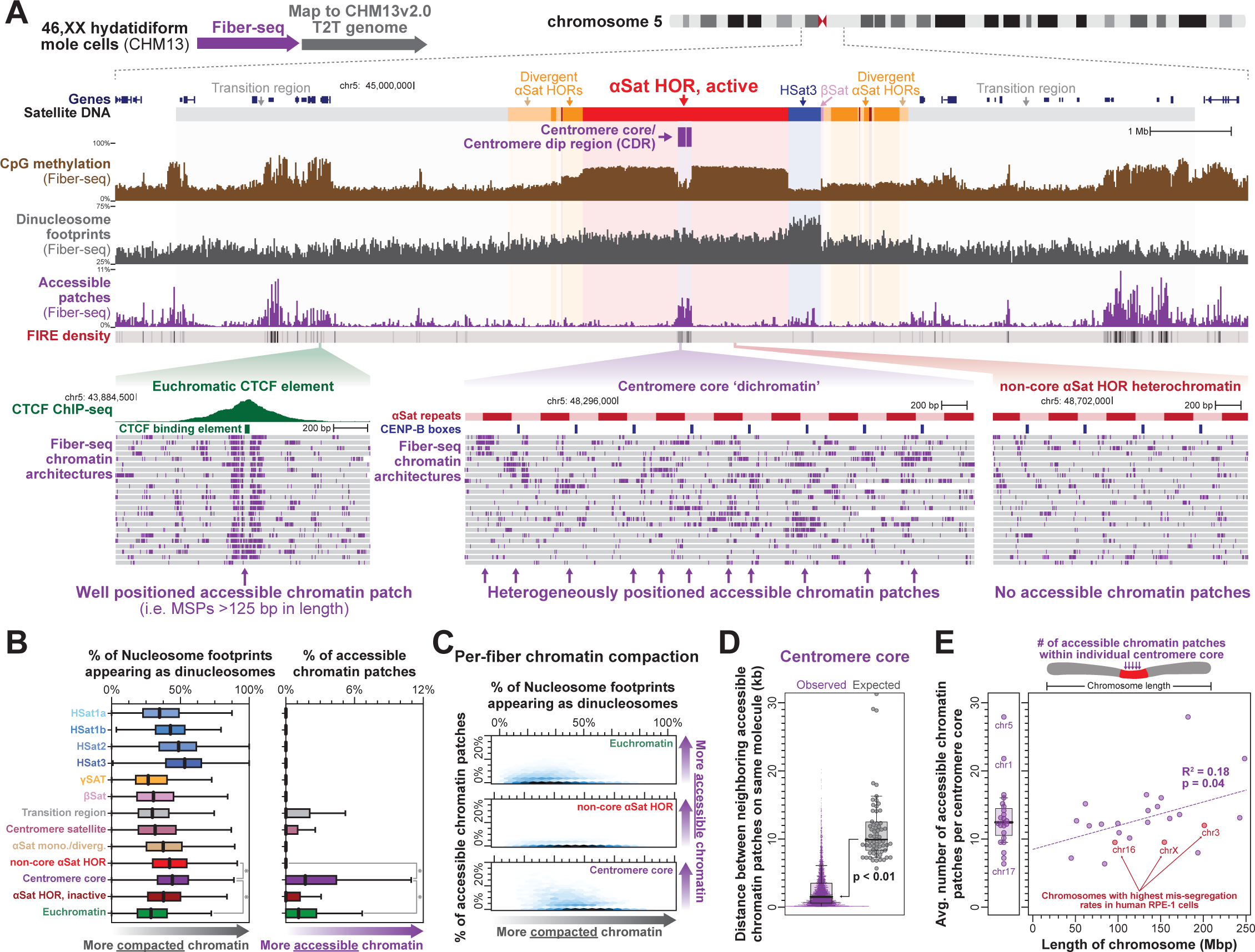
Dichotomous accessible chromatin patches mark human CHM13 centromere cores. (A) Genomic locus of chromosome 5 centromere showing satellite repeats, bulk CpG methylation, Fiber-seq identified di-nucleosome footprint density, Fiber-seq identified accessible chromatin patch density, and the density of Fiber-seq Inferred Regulatory Elements (FIRE). Below are individual Fiber-seq reads (grey bars) with m6A-modified bases in purple delineating single-molecule chromatin architectures within euchromatic and heterochromatic regions. (B) Average density of di-nucleosome footprints and accessible chromatin patches within various genomic regions (* p-value <0.01 Mann-Whitney). (C) Hexbin plot showing the single-molecule density of di-nucleosome footprints and accessible chromatin patches within various genomic regions. (D) Swarm and box-and-whisker plots showing the distance between accessible chromatin patches along the same molecule of DNA within the centromere core, as well as the expected distance based on the density of accessible chromatin patches within each chromosome’s centromere core (* p-value <0.01 Mann-Whitney). (E) Estimated total number of accessible chromatin patches along each chromosome’s centromere core versus the length of that chromosome. Chromosomes with high rates of mis-segregation^35^ are in red. See also Figure S1.

Notably, similar to yeast point centromeres, regional centromere cores in humans were also markedly enriched for large MTase-sensitive patches of chromatin, forming some of the most accessible chromatin domains within the entire human genome (**Figures 2B and 2C**). However, in stark contrast to traditional heterochromatin or euchromatin domains, individual chromatin fibers originating from the centromere core contained both tightly compacted nucleosome arrays and highly accessible chromatin features abutting each other (**Figures 2A and 2C**).

Accessible chromatin patches were infrequently observed within alpha-satellite regions outside of the centromere core, indicating that they are a unique feature of kinetochore binding. Although these accessible chromatin patches were heterogeneously placed across individual chromatin fibers within the centromere core, these patches nonetheless clustered along individual chromatin fibers within the centromere core (**Figure 2D**). In addition, each chromosome molecule containing only 6-28 accessible chromatin patches within its 80-220 kbp centromere core (**Figure 2E**), a value that mirrors estimates of the number of kinetochore microtubules that form on individual human centromeres^40,41^. Notably, longer chromosomes have significantly more accessible chromatin patches than shorter chromosomes, and after controlling for chromosome length, we observed that chromosomes with the highest rate of missegregation in human RPE-1 cells^42^ had among the lowest amount of accessible chromatin patches (**Figure 2E**). Together, these findings establish that the centromere core in human CHM13 cells contains a dichotomous chromatin organization (i.e. dichromatin) not found elsewhere in the genome, which is characterized by highly accessible chromatin patches that likely directly relate to kinetochore attachment and function within the centromere core.

### CENP-B guides centromere core dichromatin to mirror the alpha-satellite DNA repeat

We next sought to investigate the impact that alpha-satellite repeat sequence has on dichromatin formation within CHM13 centromere cores. The 170 bp alpha-satellite repeat is known to preferentially position CENP-A containing nucleosomes both *in vitro* and *in vivo*^38,43^, so we first sought to identify whether we could observe this preferential positioning along extended alpha-satellite arrays using Fiber-seq data. To identify the most common represented spacing of m6A-modified bases (i.e., the chromatin/nucleosome repeat unit length), we applied a Fourier Transform to m6A-modified bases present along each fiber. We observed that chromatin fibers originating from euchromatic and non-alpha satellite heterochromatic genomic regions have nucleosomes repeating every ∼179-190 bp (**Figures 3A, 3B and S2**), consistent with prior reports^44,45^. In contrast, centromere core alpha-satellite regions contained a nucleosome repeat unit of 170 bp, mirroring the underlying repeat length of alpha-satellite DNA (**Figure 3A and 3B**). This reveals that the predominant positioning of nucleosome arrays within the centromere core is directly related to the size of the underlying alpha-satellite repeat. However, inactive alpha-satellite HORs as well as divergent/monomeric alpha-satellite HORs contained a nucleosome repeat unit of 190-193 bp (**Figure 3B**), indicating that the alpha-satellite repeat in and of itself is not sufficient for enabling chromatin to mirror the underlying DNA repeat.

**Figure 3.**
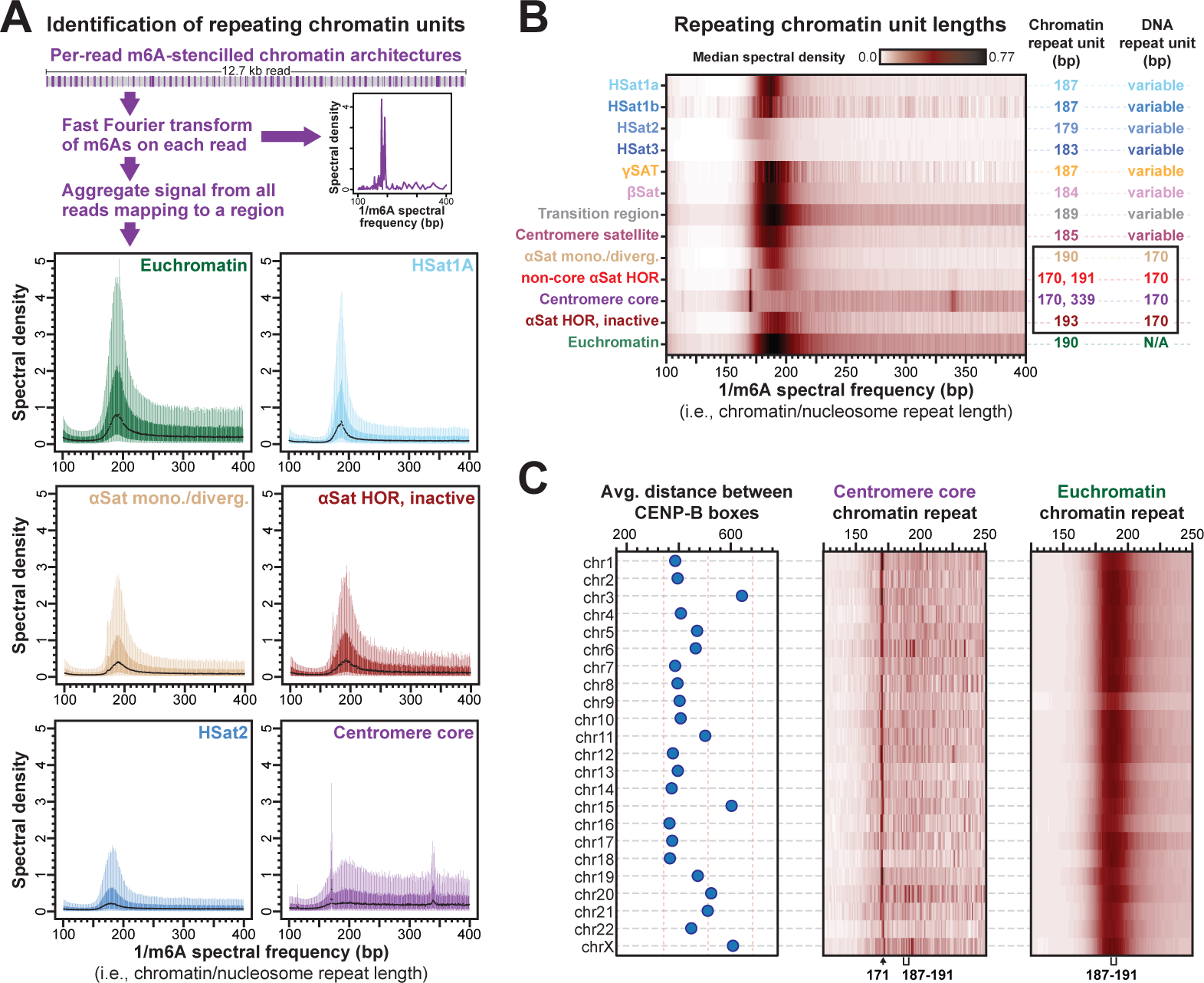
Centromere core chromatin mirrors the alpha-satellite DNA repeat. (A) Box-and-whisker plots of the chromatin repeat lengths from various genomic regions. Specifically, m6A-marked chromatin features from individual chromatin fibers were subjected to Fourier transform and the per-molecule spectral densities were then aggregated across different fibers from the same genomic region. (B) Heatmap of the median spectral density for various chromatin repeat lengths, and the peak chromatin repeat length(s) from fibers contained within the euchromatic genome, or various satellite repeat regions. The satellite DNA repeat unit for each region is also indicated. (C) Plot showing the average distance between CENP-B boxes along each centromere core within CHM13 cells, alongside a heatmap of the median spectral density for the centromere core and euchromatic regions along each centromere. See also Figure S2.

In addition to CENP-A, alpha-satellite repeats within centromere cores in humans are also bound by the sequence-specific DNA binding protein CENP-B, which occupies a non-palindromic A/T-rich 17 bp CENP-B box^3,4^ that predominantly punctuates alternating repeats within alpha-satellite HORs^46^ (**Figures 3C and S3A**). CENP-B occupancy is known to induce neighboring nucleosome positioning along these repeats *in vitro*^47,48^, and CpG methylation is a known modifier of CENP-B affinity and occupancy *in vitro*^49,50^. Given these features, we next sought to determine whether CENP-B occupancy could be playing a role in the phasing of dichromatin to the alpha-satellite repeat within the centromere core. CENP-B occupancy along the CENP-B box is readily resolved *in vitro* using DNaseI footprinting^47^, and we similarly found that CENP-B occupancy can be resolved *in vivo* using Fiber-seq (**Figures 4A and 4B**). Specifically, CENP-B boxes located within the centromere core demonstrate occupancy with an MTase-footprint that mirrors the *in vitro* CENP-B DNaseI footprint and the protein-DNA contacts within the crystal structure of the CENP-B-DNA complex^3,47,51^ (**Figures 4B**). Furthermore, consistent with prior *in vitro* findings^49,50^, we observed that CpG methylation at even one of the 2 CpG dinucleotides within the CENP-B box largely abrogates CENP-B occupancy within the centromere core *in vivo* (**Figures 4B, 4C and S3B**), indicating that CpG methylation plays a dominant role in modulating CENP-B occupancy within the centromere core. Notably, we found that CENP-B boxes that lack CpG methylation within the centromere core preferentially overlap accessible chromatin patches and position nucleosomes immediately adjacent to them (**Figures S3C and S3D**), thereby organizing dichromatin within the centromere core relative to the underlying alpha-satellite DNA repeat. However, this feature was limited to the centromere core, as CENP-B boxes outside of the centromere core were largely unoccupied irrespective of their CpG methylation status (**Figure 4C**), with weak CENP-B occupancy only observed at sporadic unmethylated CENP-B boxes located within alpha-satellite HORs flanking the centromere core, a pattern consistent with prior imaging studies^7^. Together, these findings suggest a model whereby CENP-B occupancy at hypo-CpG methylated CENP-B boxes within the centromere core is organizing chromatin architecture to mirror the underlying alpha-satellite repeat.

**Figure 4.**
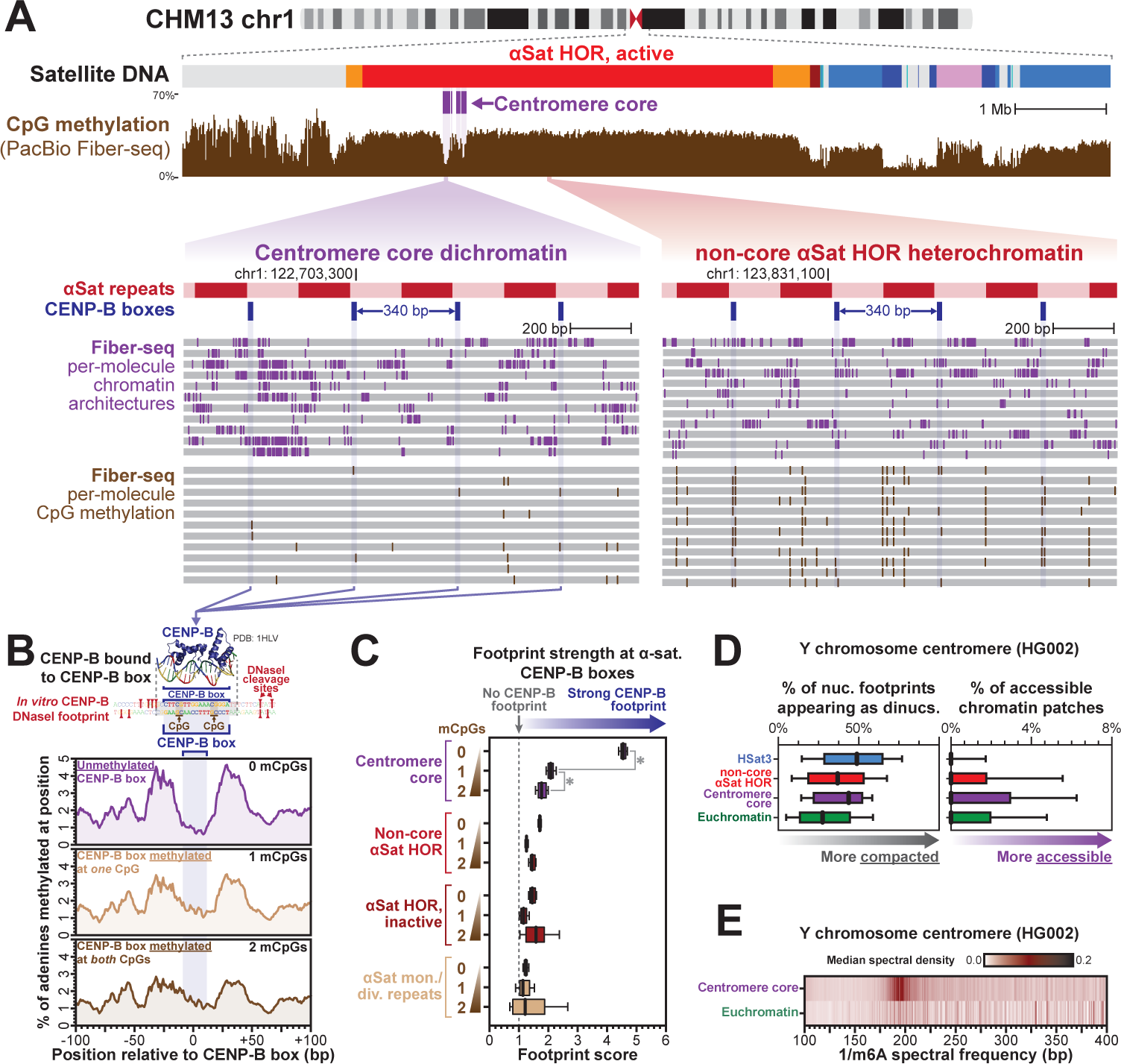
CENP-B selectively occupies and phases dichromatin within the centromere core. (A) Genomic locus showing bulk CpG methylation and per-molecule m6A-marked chromatin architectures and CpG methylations demonstrating single-molecule footprints at centromere core CENP-B boxes. (B) Structure of CENP-B bound to a CENP-box (PDB 1HLV), as well as in vitro DNaseI footprint of a CENP-B-occupied CENP-B box relative to aggregate m6A methylation profile at centromere core CENP-B boxes containing various levels of methylated CpGs (mCpGs). (C) Box-and-whisker plots of footprint scores at CENP-B boxes within various satellite regions as a function of mCpG status at the CENP-B box. Higher scores quantitatively indicate greater CENP-B occupancy. (D) Average density of di-nucleosome footprints and accessible chromatin patches within various genomic regions along the Y chromosome from HG002 using Fiber-seq data from the GM24385 cell line. (E) Heatmap of the median spectral density for centromere core and euchromatic regions along the Y chromosome from HG002 using Fiber-seq data from the GM24385 cell line. See also Figure S3.

To disentangle the role of CENP-B in dichromatin formation, we evaluated chromatin architectures along the Y chromosome centromere, which is formed from alpha-satellite repeats that lack CENP-B boxes^52^. Application of Fiber-seq to 46,XY GM24385 cells followed by mapping to the complete GM24385 reference genome (*i.e.,* HG002)^10^ demonstrated a marked enrichment in both compacted di-nucleosome footprints, as well as large patches of accessible chromatin within the Y-chromosome centromere core (**Figures 4D and S3E**). However, in contrast to studies using short-read sequencing method^38^, higher order chromatin organization within the centromere core region of chromosome Y did not mirror the underlying alpha-satellite repeat unit, but rather was a continuation of the canonical chromatin repeat unit observed throughout the rest of the HG002 genome (**Figure 4E**). Together, these findings indicate that the phasing of chromatin architectures along the alpha satellite HOR is intricately intertwined with that of CENP-B occupancy, and that dichromatin formation within these regions can form irrespective of CENP-B.

### Centromere core dichromatin is conserved across highly divergent regional centromeres

We next sought to evaluate the conservation of centromere core dichromatin architecture between multiple human genomes, as the underlying DNA sequence and structure of each centromere can markedly differ between individuals^36^. To accomplish this, we applied Fiber-seq to CHM1 and GM24385 cells, as a reference genome for all centromeres within both of these cells was recently resolved^36^. Notably, centromere core regions (*i.e.,* hypo-CpG methylated regions) within each centromere markedly diverge in both their sequence content and location when compared to CHM13 cells (**Figures 5A, 5B and S4**). However, despite the sequence divergence of centromere cores within CHM13 cells, or between CHM13 cells and CHM1 and GM24385 cells, these centromere core regions are still marked by dichromatin – exhibiting both compacted di-nucleosome footprints as well as large patches of accessible chromatin intermingled along the same chromatin fiber (**Figures 5B, 5C and S4A**). In addition, the higher order chromatin organization along the autosomes and X chromosome within CHM1 and GM24385 cells (*i.e.* centromeres that contain CENP-B boxes) similarly displayed a chromatin repeat unit of 170 bp selectively within centromere cores (**Figure S4B and S4C**). Together, these findings demonstrate that dichromatin is a conserved feature of chromatin compaction within the centromere core in humans.

**Figure 5.**
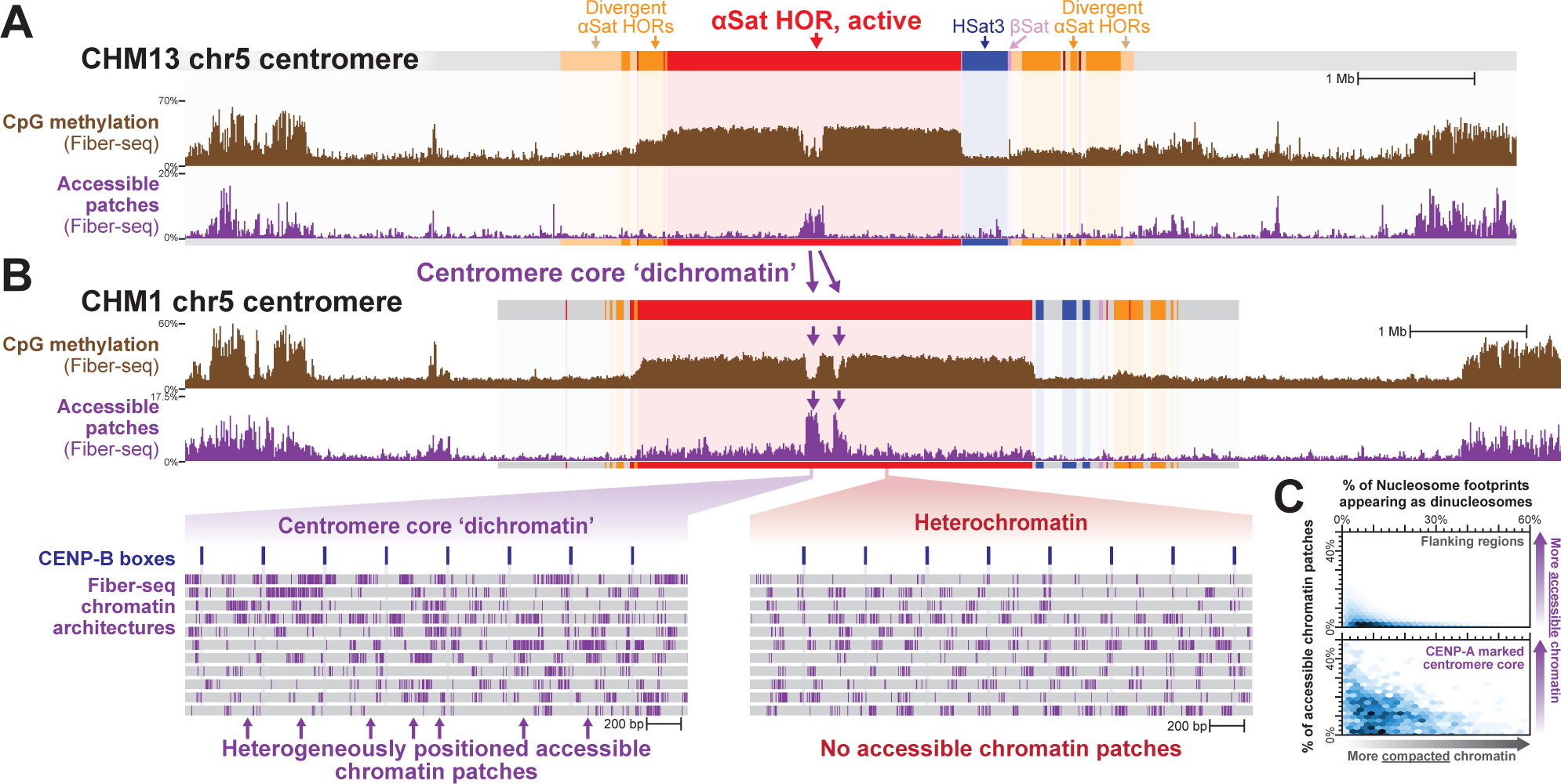
Dichromatin is a conserved feature of centromere core chromatin in humans. (A-B) Genomic locus showing chromosome 5 centromere in both CHM13 (A) and CHM1 (B) cells, including CpG methylation and accessible chromatin patches. (bottom) Single-molecule chromatin architecture along individual alpha-satellite repeats within centromere core and non-core alpha-satellite repeats. (C) Hexbin plot showing the single-molecule density of di-nucleosome footprints and accessible chromatin patches within various CHM1 centromere cores and surrounding regions. See also Figure S4.

To understand if dichromatin is a feature of regional centromeres that lack alpha-satellite repeats, we next applied Fiber-seq to resolve chromatin architectures within gibbon centromeres. Gibbons have undergone rapid karyotypic evolution accompanied by the inactivation of many ancestral centromeres and the formation of evolutionary new centromeres^53^ that are largely composed of transposable elements that lack CENP-B boxes and alpha-satellite repeats^54^. We first assembled and validated the sequence of five centromeres from the eastern hoolock gibbon (*Hoolock leuconedys*) by applying Oxford Nanopore (ONT) sequencing, Hi-C, Fiber-seq, and CENP-A CUT&RUN to a 38,XX lymphoblastoid cell line derived from Betty (**Figure 6A**). These centromeres contained hypo-CpG methylated regions corresponding to sites of CENP-A occupancy (**Figure 6B and S5**). However, the sequence of these centromere cores was largely derived from transposable elements, not alpha satellite repeats. Notably, despite the lack of alpha-satellite repeats within these centromere cores, these regions were still marked by dichromatin – exhibiting both compacted di-nucleosome footprints as well as clustered patches of accessible chromatin intermingled along the same chromatin fibers (**Figure 6B, 6C and S5E**). Unlike human autosomes, the higher order chromatin organization within the eastern hoolock gibbon centromere core regions appeared to be a continuation of the canonical chromatin repeat unit observed within flanking regions, albeit less organized owing to the dichromatin features (**Figure 6D**). Together, these findings demonstrate that dichromatin is a conserved feature of centromere core chromatin compaction irrespective of the underlying DNA sequence.

**Figure 6.**
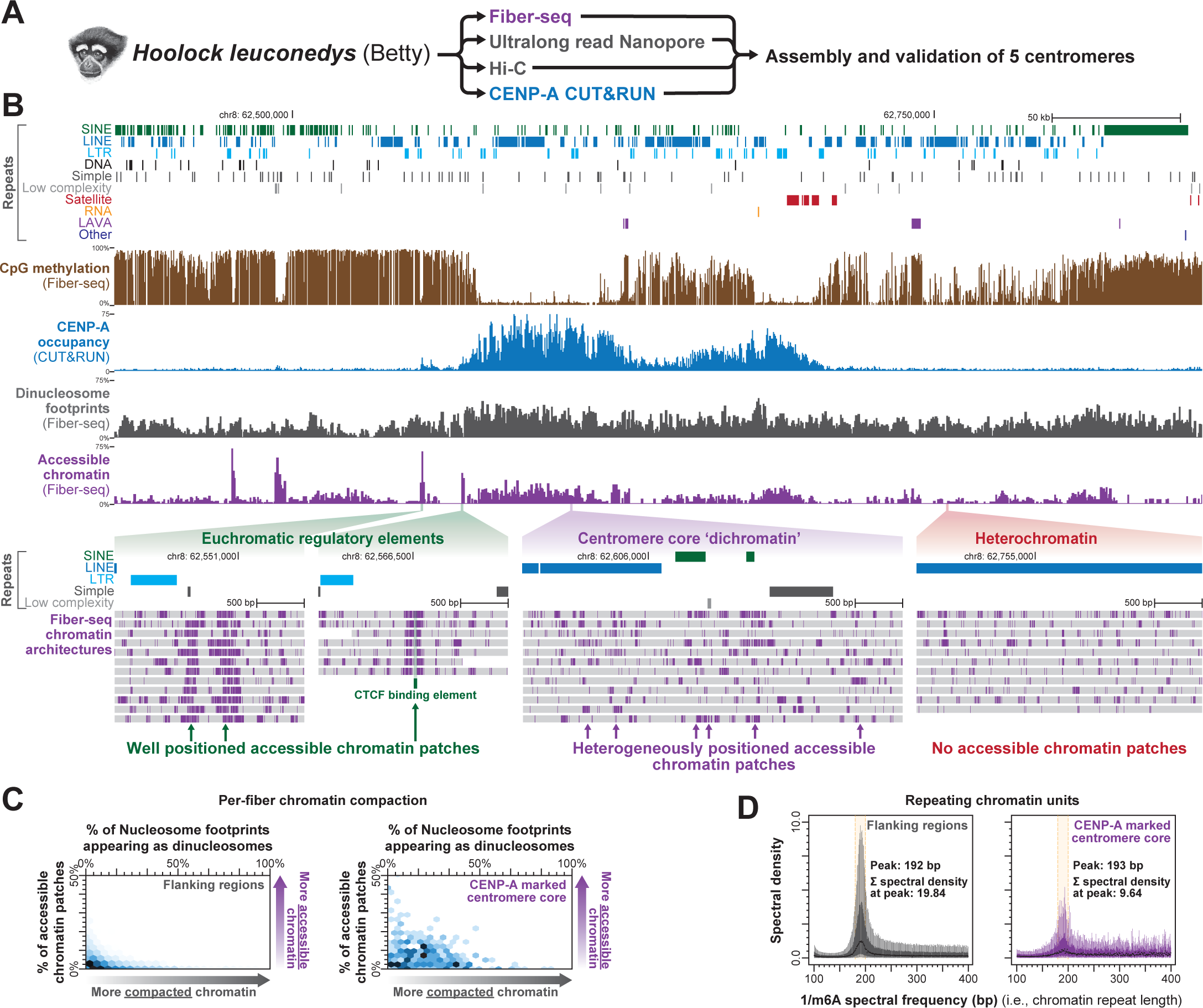
Centromere core dichromatin does not require alpha-satellite repeats. (A) Schematic for Fiber-seq in a lymphoblastoid cell line from the eastern hoolock gibbon (*Hoolock leuconedys*) Betty. Ultralong read Nanopore sequencing was used for *de novo* genome assembly, which was validated using the contigs assembled using Fiber-seq sequencing data. CENP-A CUT&RUN was used to identify centromere cores along validated assembly regions. (B) Genomic locus showing chromosome 8 centromere, including different repeat classes, CpG methylation, CENP-A CUT&RUN, and Fiber-seq derived di-nucleosome footprint density, and chromatin accessibility. (bottom) Single-molecule chromatin architecture from pericentromeric gene regulatory elements, centromere core, and heterochromatin. (C) Hexbin plot showing the single-molecule density of di-nucleosome footprints and accessible chromatin patches within centromere core and flanking regions. (D) Box-and-whisker plots of the chromatin repeat lengths from centromere core and flanking regions. Below are indicated the peak chromatin repeat unit as well as the medial spectral density at that repeat unit showing that chromatin is more disorganized within the centromere core. See also Figure S5.

## Discussion

We demonstrate that single-molecule chromatin fiber sequencing (Fiber-seq) enables the delineation of chromatin architectures within point and regional centromeres at single-molecule and single-nucleotide resolution. Application of Fiber-seq to point centromeres in yeast, as well as regional centromeres in four human and non-human primate samples with complete centromere assemblies exposed a unique form of chromatin compaction within regional centromere cores that is not observed elsewhere along the genome. Specifically, regional centromere cores simultaneously contain both the most accessible and compacted chromatin in the genome, with these dichotomous chromatin features both present along individual chromatin fibers. In addition, unlike traditional euchromatic regulatory elements, accessible chromatin patches within the centromere core are heterogeneously positioned across individual chromatin fibers. These dichotomous features of chromatin compaction within the centromere core diverge from the standard model of heterochromatin and euchromatin compaction and is hence termed ‘dichromatin’.

These accessible chromatin patches that punctuate the centromere core of regional centromeres are also a feature of point centromeres, where they create a highly plastic chromatin topology at the site of kinetochore attachment. Within regional centromeres, clustered accessible chromatin patches disrupt the ability of compacted nucleosome arrays to extend beyond ∼1 kb in length, in stark contrast to the chromatin surrounding regional centromeres core, which is marked by tightly compacted nucleosome arrays that often stretch for >20 kb in length^17^.

Centromeres comprise the most rapidly evolving DNA sequences in eukaryotic genomes^59^, and the marked heterogeneity of chromatin organization within the centromere core likely enables kinetochore attachment to be particularly immune to sequence and structural variability within centromeres. Consistent with this, we find that the dichromatin architecture of centromere cores is highly conserved between individuals, despite the marked divergence of the underlying alpha-satellite organization. Furthermore, dichromatin can assemble within centromere cores that lack alpha-satellite repeats, indicating that functional conservation within centromeres is mediated at the level of chromatin, not DNA. These findings are consistent with the centromere paradox^58^, and raise additional questions as to how the centromere core remains faithfully localized to a specific region along the genome despite its predilection to form along divergent sequences.

## Acknowledgements

We thank Steven Henikoff, Stirling Churchman and John Stamatoyannopoulos for their helpful comments and feedback. We are grateful to Nam Pho and the research computing group at the University of Washington for computational assistance. We thank the Telomere-to-Telomere (T2T) consortium for generating a complete reference genome of CHM13 cells, as well as the Y chromosome of HG002. This research was supported by NIH grant 1DP5OD029630 to A.B.S. and a 2021 Catalytic Collaborations pilot grant from the Brotman Baty Institute for Precision Medicine. A.B.S. holds a Career Award for Medical Scientists from the Burroughs Wellcome Fund and is a Pew Biomedical Scholar. S.B. and S.H. were supported by NIH R35 GM149357 and S.B. is also an investigator of the Howard Hughes Medical Institute.

## Author contributions

A.B.S., J.R. G.A.H., Y.M., S.H., K.M.M., and B.M. designed and performed the experiments. A.B.S., D.D., A.E.S.C., G.A.H., G.A.L., E.E.E., R.J.O and T.R. performed the computational analyses. A.B.S. and D.D. wrote the manuscript.

## Declaration of interests

A.B.S. is a co-inventor on a patent relating to the Fiber-seq method.

## Data and code availability

All data used for this paper will be available at GEO. All code used for this paper will be available at a Zenodo repository.

## Inclusion and diversity statement

One or more of the authors of this paper self identifies as an underrepresented ethnic minority in their field of research or within their geographical location.

## STAR METHODS

### Single-molecule chromatin fiber sequencing

CHM13 cells were grown in AmnioMax C-100 Basal Medium (Invitrogen 12558011), which includes the AmnioMAX™ C-100 Supplement to approximately 90% confluency at 37°C and 5% CO_2_. Growth media was changed daily, and cells were split using 0.25% trypsin (GIbco 25200056). One million CHM13 cells per sample were pelleted at 250 x g for 5 minutes (4 samples were processed in parallel), and washed once with PBS and then pelleted again at 250 x g for 5 minutes. Each cell pellet was resuspended in 60 μl Buffer A (15 mM Tris, pH 8.0; 15 mM NaCl; 60 mM KCl; 1mM EDTA, pH 8.0; 0.5 mM EGTA, pH 8.0; 0.5 mM Spermidine) and 60 μl of cold 2X Lysis buffer (0.1% IGEPAL CA-630 in Buffer A) was added and mixed by gentle flicking then kept on ice for 10 minutes. Samples were then pelleted at 4°C for 5 min at 350 x g and the supernatant was removed. The nuclei pellets were gently resuspended individually with wide bore pipette tips in 57.5 μl Buffer A and moved to a 25°C thermocycler. 1 μl of Hia5 MTase (200 U) ^19^ and 1.5 μl 32 mM S-adenosylmethionine (NEB B9003S) (0.8 mM final concentration) were added, then carefully mixed by pipetting the volume up and down 10 times with wide bore tips. The reactions were incubated for 10 minutes at 25°C then stopped with 3 μl of 20% SDS (1% final concentration) and transferred to new 1.5 mL microfuge tubes. The sample volumes were adjusted to 100 μl by the addition of 37 μl PBS. To that, 500 μl of HMW Lysis Buffer A (Promega Wizard HMW DNA Extraction Kit A2920) and 3 μl RNAse A (ThermoFisher EN0531) were added. The tubes were mixed by inverting 7 times and incubated 15 minutes at 37°C. 20 μl of Proteinase K (Promega Wizard HMW DNA Extraction Kit A2920) was added and the samples mixed by inverting 10 times and incubated 15 minutes at 56°C followed by chilling on ice for 1 minute. Protein was precipitated by the addition of 200 μl Protein Precipitation Solution (Promega Wizard HMW DNA Extraction Kit A2920). Using a wide bore tip the samples were mixed by drawing up contents from the bottom of the tube and then expelled on the side of the tube 5 times. Tubes were centrifuged at 16,000g for 5 minutes. The supernatant was poured into a new 1.5 ml tube containing 600 μl isopropanol. This was mixed by gentle inversion 10 times, incubated 1 minute at room temperature then inverted an additional 10 times. DNA was precipitated by centrifugation at 16,000g for 5 minutes. The supernatant was decanted and the pellet washed with the addition of 70% ethanol and centrifuged again at 16,000 g 5 minutes. After the supernatant was decanted, a quick spin was performed to facilitate removal of any residual ethanol. Open tubes were air dried on the bench for 15 minutes. The DNA was resuspended by the addition of 25 μl 10mM Tris pH 8.0. Tubes were stored overnight in 4°C and the next day were mixed gently by using a wide bore pipette tip, drawing up the sample 5 times and gently expelling the contents on the side of the tube. Samples were then stored at –80°C prior to library construction.

DNA shearing, library construction, and PacBio Sequel II sequencing were performed as previously described^20^ with the exception that 15-20 kb fragments were targeted when shearing using the Megaruptor (Diagenode Diagnostics). In addition, we performed a high-pass size selection of the SMRTbell library using the Sage Science PippinHT platform (Sage Science cat. no. ELF0001) according to the manufacturer’s protocol, using a high-pass cutoff of 10-15 kb to target an average library size of 17-20 kb.

Fiber-seq on CHM1 cells was performed in a similar manner, and CHM1 cells were similarly grown in AmnioMax C-100 Basal Medium (Invitrogen 12558011), which includes the AmnioMAX™ C-100 Supplement. Fiber-seq on GM24385 and HLE cells were performed in a similar manner with the exception that cells were grown in the following conditions. GM24385 cells were grown in RPMI-1640 supplemented with 2mM L-Glutamine, 15% fetal bovine serum (FBS) and 1% Penicillin/Streptomycin. HLE cells were grown in RPMI-1640 supplemented with 10% FBS, 1% Pen/Strep, 1% l-glutamine, 1% non-essential amino acids, and 1% sodium pyruvate.

### Yeast strains, yeast Fiber-seq and gDNA extraction

WT *Saccharomyces cerevisiae* cells (SBY3, W303 background) were grown in 10 mL of YPD media until mid-log phase and harvested by centrifugation at 3000 x g for 5 minutes. Cells were washed once with cold dH2O and resuspended in cold KPO4/Sorbitol buffer (1 M Sorbitol; 50 mM Potassium phosphate, pH 7.5; 5 mM EDTA, pH 8.0) supplemented with 0.167% β-Mercaptoethanol. Cells were spheroplasted by addition of 0.15 µg/mL final concentration of Zymolyase T100 (Amsbio 120493-1) and incubated at 23 °C for 15 minutes on a roller drum. Spheroplasts were pelleted at 300 x g for 8 minutes at 4 °C, washed twice with cold 1 M Sorbitol, and resuspended in 58 µL of Buffer A (1 M Sorbitol; 15 mM Tris-HCl, pH 8.0, 15 mM NaCl; 60 mM KCl; 1 mM EDTA, pH 8.0; 0.5 mM EGTA, pH 8.0; 0.5 mM Spermidine; 0.075% IGEPAL CA-630). Spheroplasts were treated with 1 µL of Hia5 MTase (200 U) and 1.5 µL of 32 mM S-adenosylmethionine (NEB B9003S) for 10 minutes at 25 °C. Reaction was stopped by addition of 3 µL of 20% SDS (1% final concentration) and high molecular weight DNA was purified using the Promega Wizard® HMW DNA extraction kit (A2920).

Yeast gDNA samples were quantified using a Qubit dsDNA HS Assay according to the manufacturer’s protocol. Samples were sheared to 10-15 kb with two passes through a Covaris gTUBE at 3200rpm for 2 minutes, or until most sample was passed, per pass in an Eppendorf 5424R centrifuge. ∼50 μL per sample was recovered and barcoded using the SMRTbell® prep kit 3.0 with the SMRTbell adapter index plate 96A, according to the protocol for each kit with adjustment for larger sample volume. Samples were sequenced by UW PacBio Sequencing Services on a Sequel II SMRT Cell 8M with a 30-hour movie.

### Mapping and identifying single-molecule chromatin Features

Unaligned subreads were first converted to HiFi reads using the *ccs –hifi-kinetics*. The subsequent unaligned bam files containing HiFi reads were first aligned to the appropriate reference genome using *pbmm2 –preset CCS.* Fiber-seq reads from CHM13 cells were aligned to CHM13 v1.1 (identical to CHM13v2 without chrY). Fiber-seq reads from CHM1 cells were aligned to chm1_cens_v21.fa. Fiber-seq reads from GM24385/HG002 were aligned to HG002v1.0.1. Fiber-seq reads from the eastern hoolock gibbon (*Hoolock leuconedys*) lymphoblastoid cell line (HLE) were aligned to a de novo genome assembly produced from that cell line as described below. Yeast reads were aligned to the sacCer3 genome.

Single-molecule m6A events were called along single molecules using *fibertools-rs predict-m6A* using the default parameters.

Single-molecule mCpG events were identified using PacBio’s suite of tools. Given the later release of *jasmine* we used *primrose(v1.3)* to process CHM13 using parameters: *--min-passes 3 –-keep-kinetics.* For the remaining datasets we used *jasmine(v2.0) –-min-passes 3.* We compared CHM13 data processed with *primrose* vs *jasmine* and findings were nearly identical. Aggregate bigWig tracks containing CpG methylation information were generated using *pb-CpG-tools (v2.3.1) –modsite-mode reference –model pileup_calling_model.v1.tflite*.

Nucleosomes and MSPs were identified with *fibertools-rs add-nucleosomes* using default parameters. By default, this process is run in the background during *fibertools-rs predict-m6A*.

Nucleosome, MSP, and m6A bigWig tracks were generated by counting total occurrences across all molecules at a bp and dividing by the coverage at that bp.

### Identifying the centromere core/centromere dip region

For CHM13, annotations of various satellite DNA were downloaded from the UCSC T2T hub (http://t2t.gi.ucsc.edu/chm13/hub/t2t-chm13-v1.0/cenSatAnnotation.bigBed), and putative kinetochore bindings regions for CHM13 were searched for within ‘hor’ satellite regions using CpG methylation data that was similarly downloaded from the UCSC T2T hub (http://t2t.gi.ucsc.edu/chm13/hub/t2t-chm13-v1.0/methyFreq.bigWig). CHM1 annotations were generated as previously described (Logsdon et al., In preparation). For CHM13 and CHM1, alpha satellite repeats less than 25 bp apart were merged together and we only kept alpha satellite stretches >= 100kb. We then made windows of 1190 bp across this region with step size of 170 bp (approximate alpha satellite repeat length). We then took the mean CpG methylation frequency across all CpG sites that fall in the window. We identified all windows that were below the 35^th^ percentile of CpG methylation window frequency, and merged neighboring windows that passed this threshold. Merged windows that were 1) at least 15 kb in length, 2) had a mean CpG methylation window frequency < 20^th^ percentile, and 3) were not located in the terminal 70kb of the alpha satellite repeat region were identified as CDRs. For HG002, CDRs were identified using the same approach, but the threshold to the terminus of the alpha satellite array was lowered to 15kb. For HLE, we used CENPA CUT&RUN enriched regions to identify the centromere core, as we did not know whether to expect this region to be hypo-CpG methylated.

### Identification of dinucleosomes and large MSPs

The cutoff for identifying dinucleosomes and large MSPs was 210 and 150, respectively. For each individual fiber, features (Nucleosomes or MSPs) were compared against their respective thresholds, and a percentage value was calculated for each individual chromatin fiber.

### Measuring Distance Between Neighboring Accessible Patches

To calculate the observed distance between large MSPs we calculated the distance between large MSP start positions. To calculate the expected distance between large MSPs in the CDR, we first calculated the number of large MSPs fully contained in the CDR and divided by the total number of sequenced bases per CDR to get a large MSP / sequenced bp value. We then multiply the resulting value by the length of the respective CDR to get the expected number of large MSPs per CDR. We divide the length of the CDR by this value to get the expected number of bp between large MSPs within the CDR.

### Determining chromatin repeat lengths

To determine the chromatin repeat length of individual chromatin fibers, the per-base m6A pattern on each fiber was converted into a binary vector (m6a=1, no-m6a=0). These vectors were then independently analyzed using the *scipy.signal.periodogram* tool. This results in spectral density estimations across a vector of frequencies, which is the inverse of the nucleosome repeat length in base-pairs. Spectral densities derived from multiple fibers originating from similar genomic loci were combined and displayed.

### Delineating CENP-B binding elements and occupancy

CENP-B boxes were identified in each annotation group using the 17-bp CENP-B box NTTCGNNNNANNCGGGN motif. Specifically, genomic regions were scanned with this motif using *fimo* (version 4.11.2) ^60^ with the *--parse-genomic-coord –-qv-thresh –-thresh 0.05 –-max-stored-scores 1000000* flags. We further filtered motifs to those which contained 2 CpG sites. To delineate the occupancy of these CENP-B boxes, we intersected CENP-B boxes per-fiber methylation calls. Specifically, the m6A methylation profile of every region along every fiber overlapping a CENP-B box was converted into a binary profile (m6a=1, no-m6a=0) and stored in an array. Based on genomic context and/or CpG methylation contexts these arrays were collapsed to form m6A frequency tracks with a rolling average of 10bp.

To calculate footprint score distributions, we took all fibers fulfilling a filtering requirement (CpG methylation status / genomic region), subsampled to 10% of the fibers 1000 times and collapsed these to generate m6A frequency tracks smoothed by a 10bp window. We used these fibers to calculate a footprint score defined by the m6A frequency in the flanking region divided by the m6A frequency in the core CENP-B box. The flanking region score was defined as the mean of the smoothed m6A content –35 – –27 bases upstream and +22 – +27 bases downstream of the CENPB box center. The CENP-B core score was defined by the mean of the smoothed m6A frequency in the –7 – +1 positions relative to the center of the CENP-B box.

### Quantifying nucleosome positioning surrounding CENP-B boxes

To visualize the association of CENP-B occupancy and surrounding nucleosome positioning we first classified every instance of a chromatin fiber overlapping a CENP-B box by its CpG methylation state (0,1 or 2 mCpGs). For each overlap, we took the closest (Relative to center of CENPB box) boundary of both the upstream and downstream nucleosome. For each position upstream or downstream of the CENP-B box we then calculated the percentage of upstream or downstream nucleosomes beginning at that position, respectively.

### *Hoolock leuconedys* Oxford Nanopore Technologies (ONT) Sequencing

Cells were collected from a transformed lymphoblastoid cell line (LCL) derived from a female *Hoolock leuconedys* individual (Betty). To isolate high molecular weight (HMW) DNA, a phenol/chloroform/isoamyl alcohol extraction was performed per standard protocols for the isolation of high molecular weight DNA. Library preparation was performed using the Ligation Sequencing Kit (LSK109). HMW DNA was sequenced on the PromethION platform from Oxford Nanopore Technologies using a PromethION R9.4.1 FLOPRO002 flow cell and basecalled using Guppy (v5.0.16).

To isolate ultra-high molecular weight (UHMW) DNA, the Circulomics Nanobind UHMW DNA Extraction for cultured cells (EXT-CLU-001) protocol was followed according to manufacturer’s instructions with the Circulomics Nanobind CBB Big DNA Kit (NB-900-001-01). Library preparation was performed using the Circulomics Nanobind Library Prep protocol for ultra-long sequencing (LBP-ULN-001) using the Circulomics Nanobind UL Library Prep Kit (NB-900-601-01) and the Oxford Nanopore Technologies Ultra-Long DNA Sequencing Kit (SQK-ULK001). UHMW DNA was sequencing on the PromethION platform using a R9.4.1 FLOPRO002 flow cell and basecalled using Guppy (v5.0.16).

### *Hoolock leuconedys* Dovetail™ Omni-C™ Sequencing

Roughly 1.5 million cells were collected from the previously described *Hoolock leuconedys* lymphoblastoid cell line and processed according to the Dovetail™ Omni-C™ Proximity Ligation Assay protocol for mammalian samples (v1.4) as written. Lysate quantification was performed using the Qubit® dsDNA HS kit for the Qubit® Fluorometer and the D5000 HS kit for the Agilent TapeStation 2200. 150 bp paired end sequencing was performed on the Illumina NextSeq 550 V2 platform to a depth of ∼274M reads.

### Hoolock leuconedys CUT&RUN Sequencing

The CUT&RUN Assay Kit (#86652) from Cell Signaling Technology® was used to assess CENP-A protein-DNA interactions with 250,000 cells per condition following manufacturer’s instructions. To assess CENPA-DNA interactions, the CENP-A monoclonal antibody (ADI-KAM-CC006-E) was used at a dilution of 1:50. Tri-methyl-histone H3 (Lys4) (C42D8) rabbit monoclonal antibody (#9751) at a dilution of 1:50 was used as a positive control; rabbit (DA1E) monoclonal antibody IgG XP® isotype control (#66362) at a dilution of 1:10 was used as a negative control. Input chromatin samples were sheared to ∼100-700 bases using a Covaris S2 sonicator prior to purification. DNA purification was performed using the Cell Signaling® DNA purification with spin columns kit (#14209). DNA concentration was assessed using the Qubit® dsDNA HS kit for the Qubit® Fluorometer and the High Sensitivity D1000 kit for the Agilent TapeStation 2200. CENP-A and Input libraries were prepared using the NEBNext Ultra II DNA Library Prep Kit for Illumina (#E7645S) and sequenced using the Illumina NovaSeq 150 bp paired end settings to a depth of ∼15M reads.

### Hoolock leuconedys genome assembly

Flye (v2.9) ^61^ was used to assemble the raw Oxford Nanopore reads using an estimated genome size of 2.9 Gb, the size of the previously assembled *Nomascus leucogenys* genome ^62^. Medaka (v1.4.3) was used for long read polishing using default settings and the r941_prom_sup_g507 model.

Publicly available Illumina WGS sequencing reads for the same individual (SAMN12702557) ^63^ were used to polish the assembly by mapping the reads with Burrow’s Wheeler Aligner (v0.7.17) using the bwa mem algorithm and processed using Samtools (v1.7). Short read error correction was performed using Pilon (v1.22) with default parameters. Haplotype redundancies and assembly artifacts based on read coverage were removed using minimap2 (v2.15) and PURGEhaplotigs (v1.0). Omni-C sequences were used to scaffold the assembly with Juicer (v1.6) ^64^ and 3D-DNA (v180922) ^65^ following the default protocols outlined by developers. Juicebox with Assembly Tools (v1.11.08) was used for manual review of the produced scaffolds. Gap filling was performed using TGS-GapCloser (v1.0.1). The gap-filled assembly was polished with Illumina reads using Pilon (v.1.22) with default parameters. The reformat.sh module of BBMap was used to impose a 3kb limit on the genome.

LASTZ (v1.04.15) was used to align the assembly to the CHM13 v1.1 genome. Resulting alignments were validated according to predicted syntenic regions and large-scale chromosome misassemblies and misorientations were manually corrected using Emboss (6.6.0). To reduce any misassemblies associated with manual curation and scaffolding, pre-scaffolded contigs (the “query”) were aligned to the curated assembly (the “reference”) and scaffolded using RagTag (v2.1.0). The resulting scaffolded assembly (built from “query” contigs) was gap filled using TGS-GapCloser (v1.0.1). The gap-filled assembly was polished with Illumina reads using Pilon (v.1.22) with default parameters.

### *Hoolock leuconedys* centromere annotation

A two-step approach was used to identify contiguous centromeres for Fiber-seq analysis. First, centromeric regions were identified by enrichment of CENP-A binding. Low quality reads (Phred score < 20, length < 50 nt) and adapters were trimmed from CENP-A and Input CUT&RUN sequences using cutadapt (v3.5) and mapped using Bowtie2 with default parameters (v2.5.0). Peaks were called using MACS (v2.1.2) using the –-keep-dup=2 –-broad –q 0.01 –g 2.8e9 flags. These peaks were labeled as putative centromere regions. Second, hifiasm was run directly on the Fiber-seq reads using default parameters. Hifiasm contigs were then mapped to the Hoolock leuconedys genome assembly generated using ONT and HiC data above. Putative centromere regions that were fully encompassed within a hifiasm contig that mapped to that region were then considered to be validated as correctly assembled centromeres, as these centromeres were independently assembled using two orthologous technologies. Five centromeres passed these filters and all five were validated by manual review of read coverage and selected for subsequent Fiber-seq analysis. Repeats in the genome were annotated with RepeatMasker (v4.1.2-p1) using the Crossmatch search engine (v1.090518) and a combined gibbon (*Hylobates* sp.) Dfam (v3.6) and Repbase (20181026) repeat library. CpG methylation was detected in the ONT reads using the Bonito (v0.3.2) –Remora (v1.1.1) basecalling pipeline with default parameters and the dna_r9.4.1_e8_sup@v3.3 model. Resulting files were converted to a bedMethyl file using modbam2bed (v0.6.2).

## QUANTIFICATION AND STATISTICAL ANALYSIS

Unless noted otherwise, significance was tested using a Mann-Whitney U test. In box plots, box extends from Q1 to Q3, and whiskers represent the 10^th^ and 90^th^ percentile.

## Supplemental information

### Supplemental Figure legends

**Figure S1.**
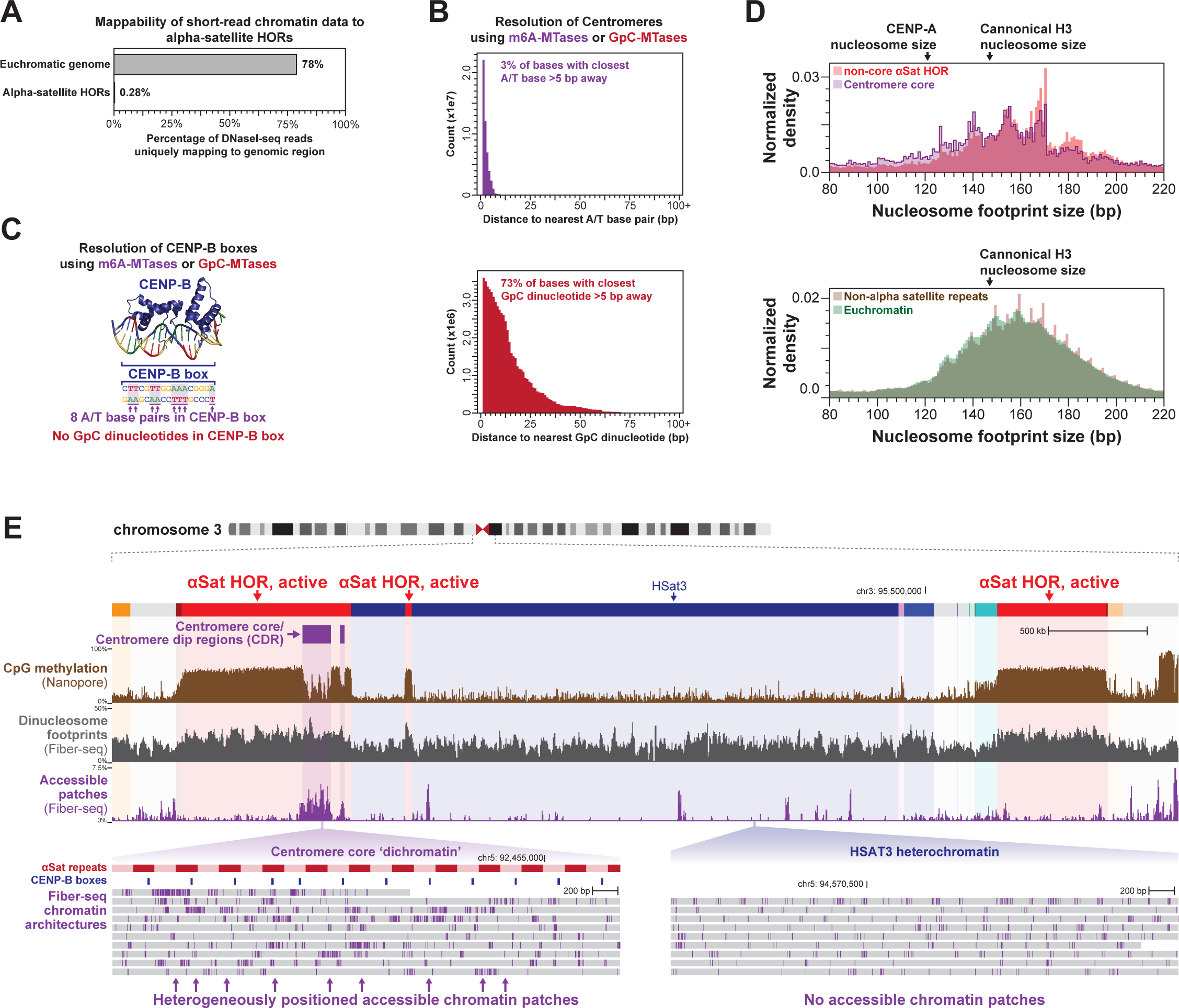
Single-molecule chromatin figure sequencing of human centromeres, related to Figure 2. (A) Bar plot showing the percentage of short-read DNaseI-seq data mapping to euchromatic genomic regions as well as alpha-satellite DNA that are uniquely mapping to these regions. (B-C) Resolution of GpC methyltransferase-based and non-specific m6A-methyltransferase-based methods for mapping chromatin architectures within the centromere (B) as well as CENP-B occupancy along CENP-B boxes (C). (D) Histogram of nucleosome footprint sizes within the centromere core, as well as in other genomic regions. (E) Genomic locus of chromosome 3 centromere showing satellite repeats, bulk CpG methylation, Fiber-seq identified di-nucleosome footprint density, as well as Fiber-seq identified chromatin accessibility density. Below are individual Fiber-seq reads (grey bars) with m6A-modified bases in purple delineating single-molecule chromatin architectures within centromere core and HSat3 heterochromatic regions.

**Figure S2.**
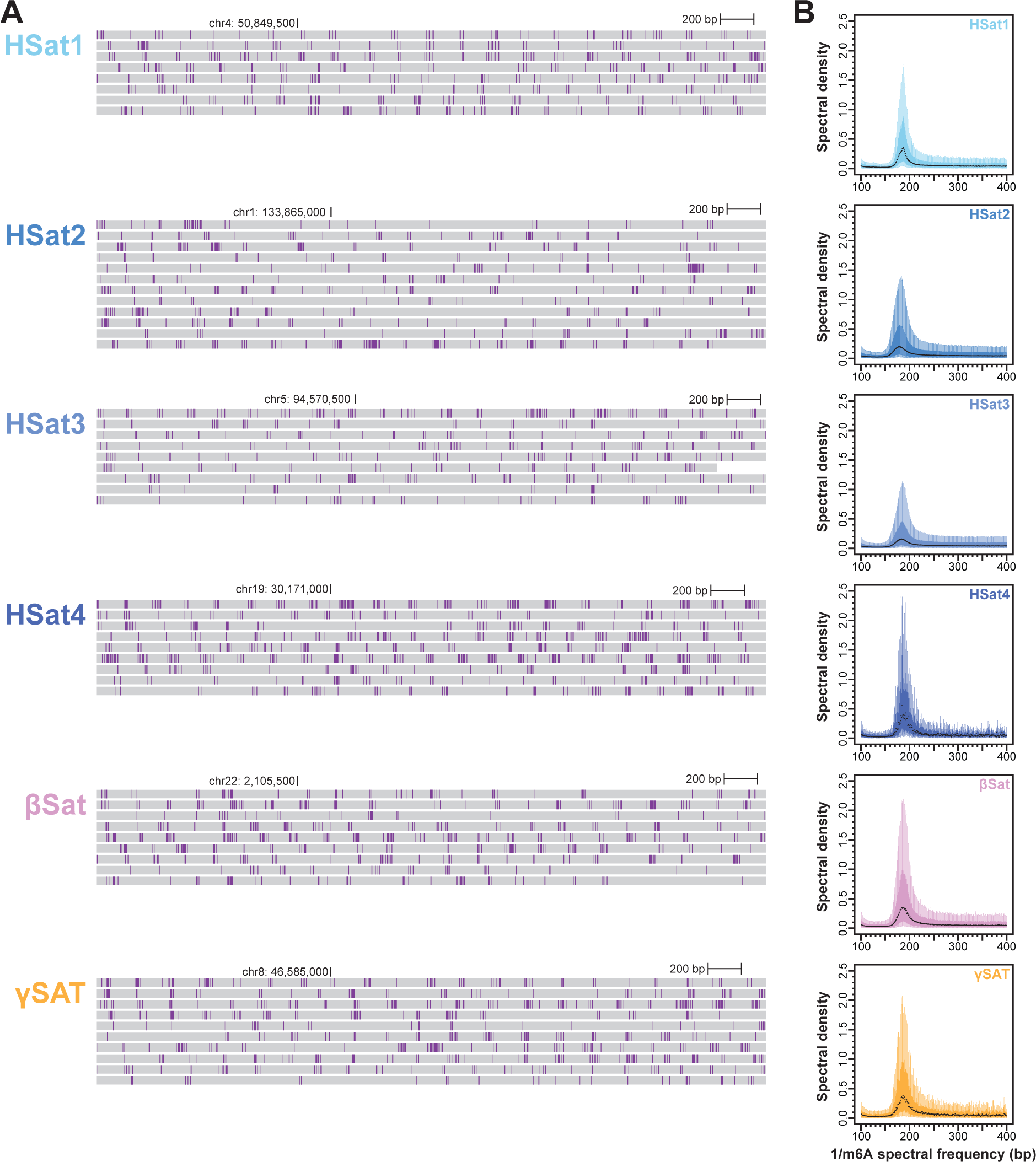
Higher-order chromatin architectures of heterochromatin, related to Figure 3. (A) Genomic loci showing single-molecule chromatin architectures within six separate satellite regions. (B) Box-and-whisker plots of the chromatin repeat lengths from each of these six distinct satellite regions.

**Figure S3.**
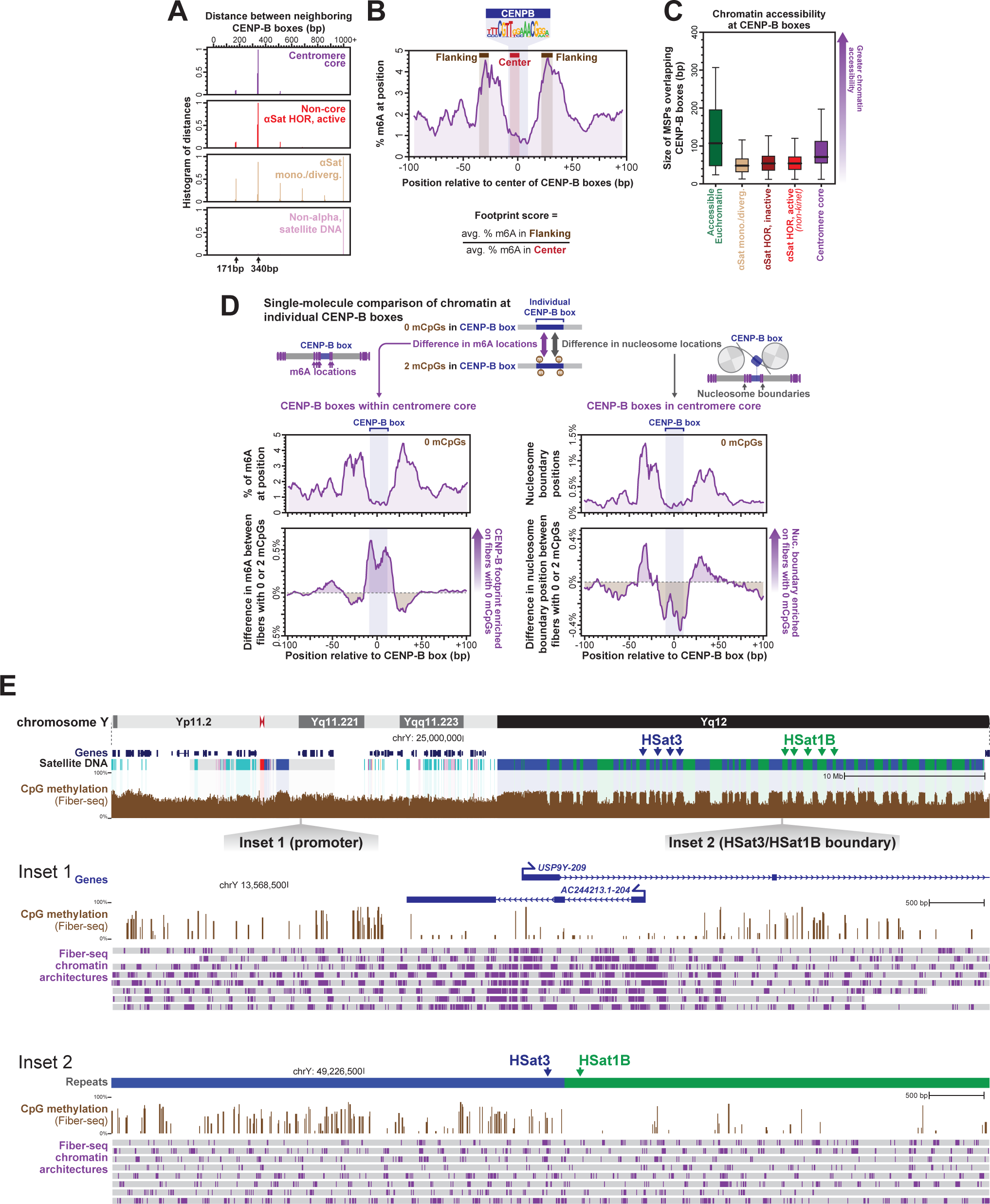
Single-molecule CENP-B occupancy within centromere core, related to Figure 4. (A) Histogram of the distance between CENP-B boxes within various genomic regions (B) Aggregate m6A methylation profile at all CENP-B boxes located within the centromere core. Regions used for calculating footprint score are highlighted, and the calculation of the footprint score is enumerated below. (C) Box-and-whisker plots showing the size of MSPs overlapping CENP-B boxes identified within different genomic regions. (D) Directly comparing m6A-accessibility and nucleosome footprint boundaries at individual CENP-B boxes based on whether the read does or does not contain mCpG within that CENP-B box. Above is the aggregate m6A methylation (left) or nucleosome boundary positions (right) at centromere core alpha satellite CENP-B boxes containing no mCpG. Below is the aggregate difference in these features between reads with and without mCpG overlapping the CENP-B box. (E) Genomic locus showing CpG methylation signal along the entire chromosome Y in HG002/GM24385 cells. Below are insets of single-molecule chromatin architectures at a promoter within the euchromatic region of chromosome Y, as well as an HSat3-HSat1B boundary within the heterochromatic q arm of chromosome Y.

**Figure S4.**
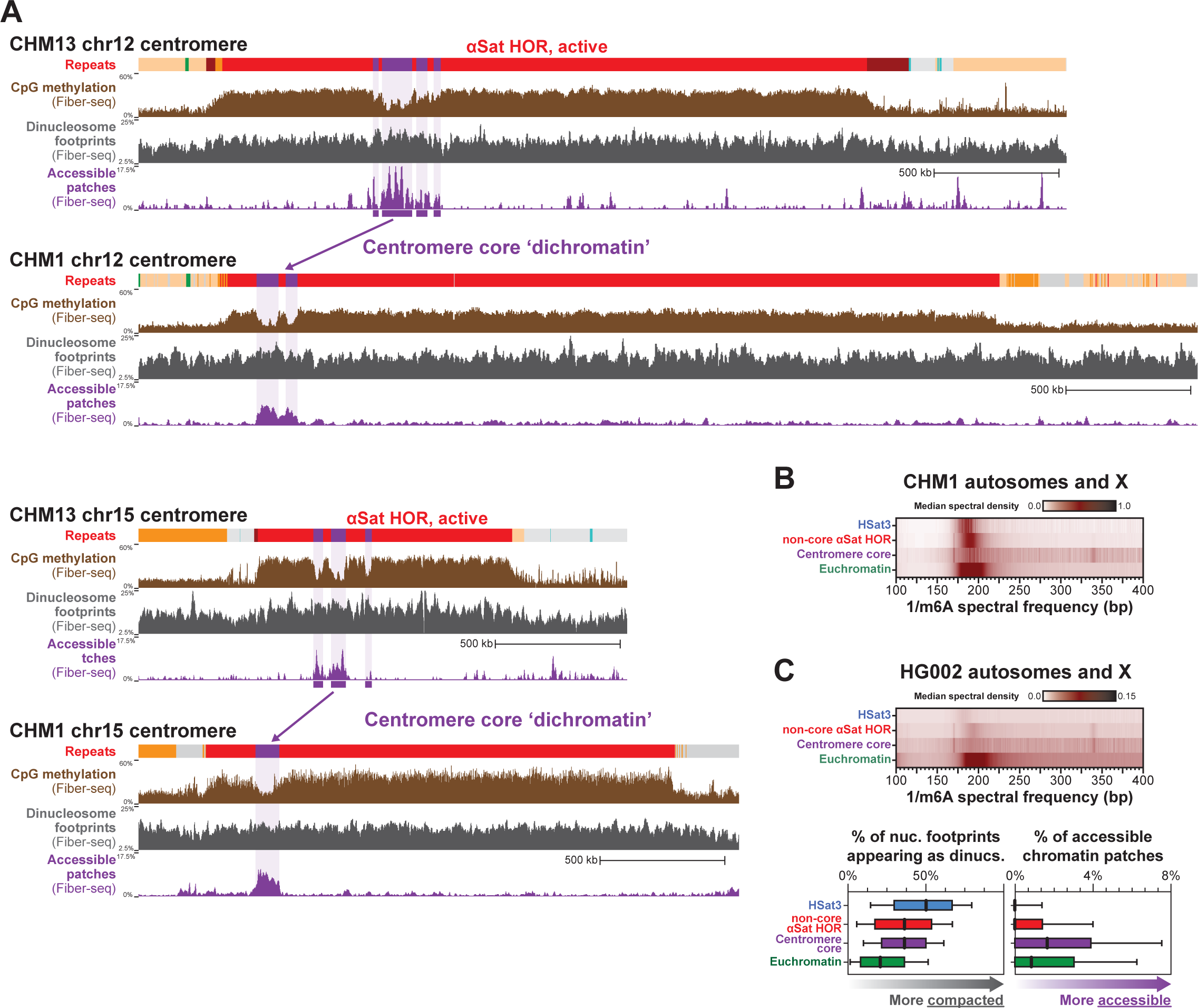
Conservation of dichromatin architecture between humans, related to Figure 5. (A) Genomic loci showing CpG methylation signal as well as di-nucleosome density and chromatin accessibility along the chromosome 12 and 15 centromeres in both CHM13 and CHM1 cells. (B) Heatmap of the median spectral density for various genomic regions along the autosomes and X chromosome from CHM1 using Fiber-seq data from the CHM1 cell line. (C) (top) Heatmap of the median spectral density for various genomic regions along the autosomes and X chromosome from HG002 using Fiber-seq data from the GM24385 cell line. (bottom) Average density of di-nucleosome footprints and accessible chromatin patches within various genomic regions along the autosomes and X chromosome from HG002 using Fiber-seq data from the GM24385 cell line.

**Figure S5.**
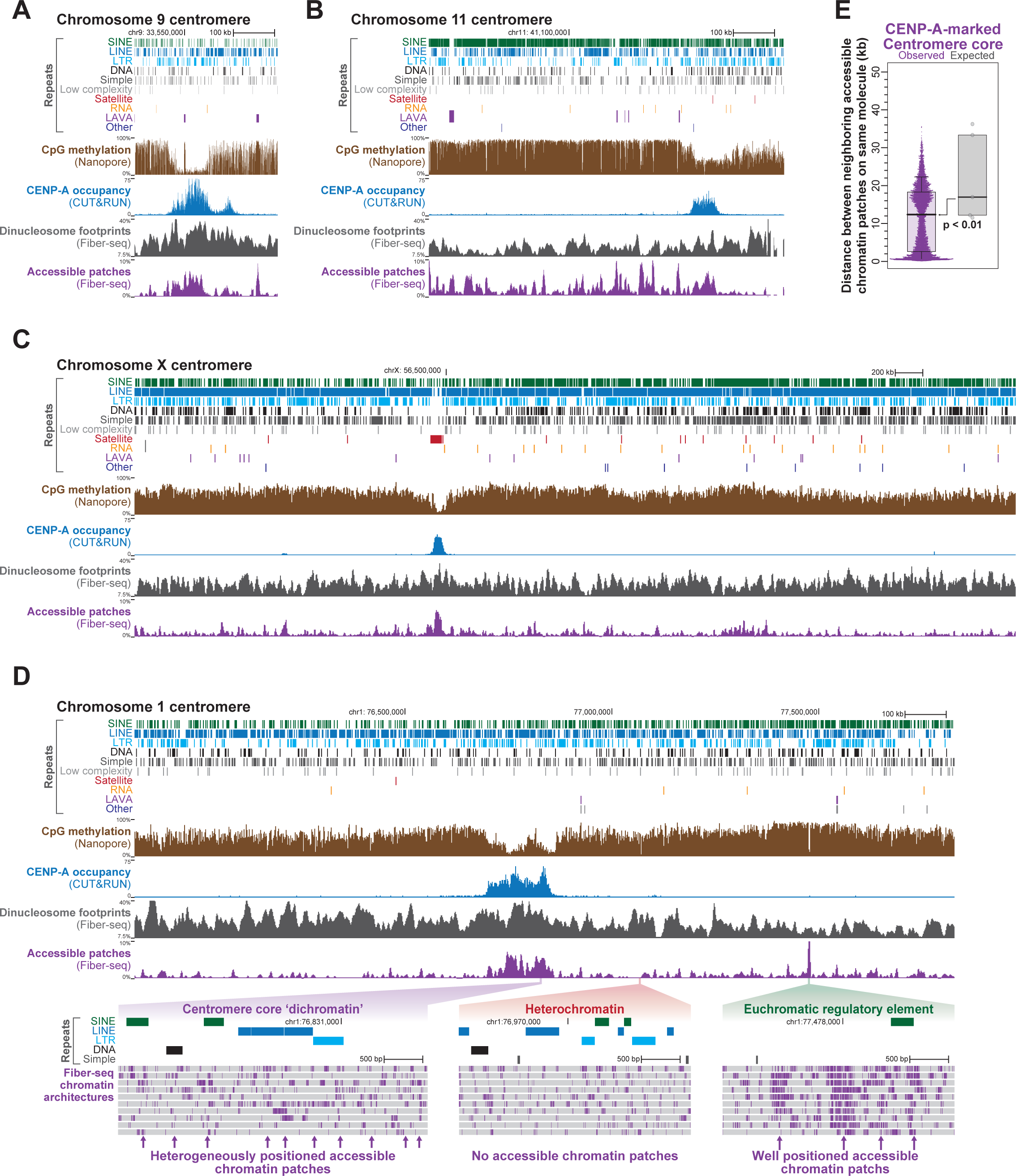
Alpha-satellite DNA is not necessary for dichromatin formation, related to Figure 6. (A-D) Genomic loci showing repeats, CpG methylation, CENP-A CUT&RUN, and Fiber-seq derived di-nucleosome density and chromatin accessibility along four assembled centromeres from a lymphoblastoid cell line from the eastern hoolock gibbon (*Hoolock leuconedys*) Betty. Below inset showing single-molecule chromatin architectures within the centromere core region, a heterochromatic region, and a gene regulatory element along chromosome 1. (F) Swarm and box-and-whisker plots showing the distance between accessible chromatin patches along the same molecule of DNA within the centromere core, as well as the expected distance based on the density of accessible chromatin patches within each chromosome’s centromere core (* p-value <0.01 Mann-Whitney).

## Notes

### Summary of Updates

There was an inadvertent error with the prior Figure 5C that has now been corrected.

## References

1. Rosenberg, H., Singer, M., and Rosenberg, M. (1978). Highly reiterated sequences of SIMIANSIMIANSIMIANSIMIANSIMIAN. Science 200, 394–402.

2. Altemose, N., Logsdon, G.A., Bzikadze, A.V., Sidhwani, P., Langley, S.A., Caldas, G.V., Hoyt, S.J., Uralsky, L., Ryabov, F.D., Shew, C.J., et al. (2022). Complete genomic and epigenetic maps of human centromeres. Science 376. 10.1126/SCIENCE.ABL4178.

3. Muro, Y., Masumoto, H., Yoda, K., Nozaki, N., Ohashi, M., and Okazaki, T. (1992). Centromere protein B assembles human centromeric α-satellite DNA at the 17-bp sequence, CENP-B box. J. Cell Biol. 116, 585–596.

4. Masumoto, H., Masukata, H., Muro, Y., Nozaki, N., and Okazaki, T. (1989). A human centromere antigen (CENP-B) interacts with a short specific sequence in alphoid DNA, a human centromeric satellite. J. Cell Biol. 109, 1963–1973.

5. Palmer, D.K., O’Day, K., Trong, H.L.E., Charbonneau, H., and Margolis, R.L. (1991). Purification of the centromere-specific protein CENP-A and demonstration that it is a distinctive histone. Proc. Natl. Acad. Sci. U. S. A. 88, 3734–3738.

6. Warburton, P.E., Cooke, C.A., Bourassa, S., Vafa, O., Sullivan, B.A., Stetten, G., Gimelli, G., Warburton, D., Tyler-Smith, C., Sullivan, K.F., et al. (1997). Immunolocalization of CENP-A suggests a distinct nucleosome structure at the inner kinetochore plate of active centromeres. Curr. Biol. 7, 901–904.

7. Fachinetti, D., Han, J.S., McMahon, M.A., Ly, P., Abdullah, A., Wong, A.J., and Cleveland, D.W. (2015). DNA Sequence-Specific Binding of CENP-B Enhances the Fidelity of Human Centromere Function. Dev. Cell 33, 314–327.

8. Zhang, W., Lee, H.R., Koo, D.H., and Jiang, J. (2008). Epigenetic modification of centromeric chromatin: Hypomethylation of DNA sequences in the CENH3-associated chromatin in Arabidopsis thaliana and maize. Plant Cell 20, 25–34.

9. Logsdon, G.A., Vollger, M.R., Hsieh, P., Mao, Y., Liskovykh, M.A., Koren, S., Nurk, S., Mercuri, L., Dishuck, P.C., Rhie, A., et al. (2021). The structure, function and evolution of a complete human chromosome 8. Nature 593, 101–107.

10. Rhie, A., Nurk, S., Cechova, M., Hoyt, S.J., Taylor, D.J., Altemose, N., Hook, P.W., Koren, S., Rautiainen, M., Alexandrov, I.A., et al. (2023). The complete sequence of a human Y chromosome. Nature 621, 344–354.

11. Sullivan, B.A., and Karpen, G.H. (2004). Centromeric chromatin exhibits a histone modification pattern that is distinct from both euchromatin and heterochromatin. Nat. Struct. Mol. Biol. 11, 1076–1083.

12. Barski, A., Cuddapah, S., Cui, K., Roh, T.-Y., Schones, D.E., Wang, Z., Wei, G., Chepelev, I., and Zhao, K. (2007). High-Resolution Profiling of Histone Methylations in the Human Genome. Cell 129, 823–837.

13. Gershman, A., Sauria, M.E.G., Guitart, X., Vollger, M.R., Hook, P.W., Hoyt, S.J., Jain, M., Shumate, A., Razaghi, R., Koren, S., et al. (2022). Epigenetic patterns in a complete human genome. Science 376, eabj5089.

14. Panchenko, T., Sorensen, T.C., Woodcock, C.L., Kan, Z.Y., Wood, S., Resch, M.G., Luger, K., Englander, S.W., Hansen, J.C., and Black, B.E. (2011). Replacement of histone H3 with CENP-A directs global nucleosome array condensation and loosening of nucleosome superhelical termini. Proc. Natl. Acad. Sci. U. S. A. 108, 16588–16593.

15. Thakur, J., and Henikoff, S. (2016). CENPT bridges adjacent CENPA nucleosomes on young human α-satellite dimmers. Genome Res. 26, 1178–1187.

16. Ando, S., Yang, H., Nozaki, N., Okazaki, T., and Yoda, K. (2002). CENP-A, –B, and –C Chromatin Complex That Contains the I-Type α-Satellite Array Constitutes the Prekinetochore in HeLa Cells. Mol. Cell. Biol. 22, 2229–2241.

17. Zhou, K., Gebala, M., Woods, D., Sundararajan, K., Edwards, G., Krzizike, D., Wereszczynski, J., Straight, A.F., and Luger, K. (2022). CENP-N promotes the compaction of centromeric chromatin. Nat. Struct. Mol. Biol. 29, 403–413.

18. Nurk, S., Koren, S., Rhie, A., Rautiainen, M., Bzikadze, A.V., Mikheenko, A., Vollger, M.R., Altemose, N., Uralsky, L., Gershman, A., et al. (2022). The complete sequence of a human genome. Science 376, 44–53.

19. Stergachis, A.B., Debo, B.M., Haugen, E., Churchman, L.S., and Stamatoyannopoulos, J.A. (2020). Single-molecule regulatory architectures captured by chromatin fiber sequencing. Science 368, 1449–1454.

20. Lee, I., Razaghi, R., Gilpatrick, T., Molnar, M., Gershman, A., Sadowski, N., Sedlazeck, F.J., Hansen, K.D., Simpson, J.T., and Timp, W. (2020). Simultaneous profiling of chromatin accessibility and methylation on human cell lines with nanopore sequencing. Nat. Methods 17, 1191–1199.

21. Shipony, Z., Marinov, G.K., Swaffer, M.P., Sinnott-Armstrong, N.A., Skotheim, J.M., Kundaje, A., and Greenleaf, W.J. (2020). Long-range single-molecule mapping of chromatin accessibility in eukaryotes. Nat. Methods, 1–9.

22. Abdulhay, N.J., McNally, C.P., Hsieh, L.J., Kasinathan, S., Keith, A., Estes, L.S., Karimzadeh, M., Underwood, J.G., Goodarzi, H., Narlikar, G.J., et al. (2020). Massively multiplex single-molecule oligonucleosome footprinting. Elife 9, 1–23.

23. Vollger, M.R., Swanson, E.G., Neph, S.J., Ranchalis, J., Munson, K.M., Ho, C.-H., Sedeño-Cortés, A.E., Fondrie, W.E., Bohaczuk, S.C., Mao, Y., et al. (2024). A haplotype-resolved view of human gene regulation. bioRxiv, 2024.06.14.599122. 10.1101/2024.06.14.599122.

24. Drozdz, M., Piekarowicz, A., Bujnicki, J.M., and Radlinska, M. (2012). Novel non-specific DNA adenine methyltransferases. Nucleic Acids Res. 40, 2119–2130.

25. Kong, Y., Cao, L., Deikus, G., Fan, Y., Mead, E.A., Lai, W., Zhang, Y., Yong, R., Sebra, R., Wang, H., et al. (2022). Critical assessment of DNA adenine methylation in eukaryotes using quantitative deconvolution. Science 375, 515–522.

26. Debo, B.M., Mallory, B.J., and Stergachis, A.B. (2023). Evaluation of N 6-methyldeoxyadenosine antibody-based genomic profiling in eukaryotes. Genome Res. 33, 427–434.

27. Jha, A., Bohaczuk, S.C., Mao, Y., Ranchalis, J., Mallory, B.J., Min, A.T., Hamm, M.O., Swanson, E., Dubocanin, D., Finkbeiner, C., et al. (2024). DNA-m6A calling and integrated long-read epigenetic and genetic analysis with fibertools. Genome Res. 10.1101/gr.279095.124.

28. Meluh, P.B., Yang, P., Glowczewski, L., Koshland, D., and Smith, M.M. (1998). Cse4p is a component of the core centromere of Saccharomyces cerevisiae. Cell 94, 607–613.

29. Joglekar, A.P., Bouck, D.C., Molk, J.N., Bloom, K.S., and Salmon, E.D. (2006). Molecular architecture of a kinetochore-microtubule attachment site. Nat. Cell Biol. 8, 581–585.

30. Cai, M., and Davis, R.W. (1990). Yeast centromere binding protein CBF1, of the helix-loop-helix protein family, is required for chromosome stability and methionine prototrophy. Cell 61, 437–446.

31. Lechner, J., and Carbon, J. (1991). A 240 kd multisubunit protein complex, CBF3, is a major component of the budding yeast centromere. Cell 64, 717–725.

32. Funk, M., Hegemann, J.H., and Philippsen, P. (1989). Chromatin digestion with restriction endonucleases reveals 150-160 bp of protected DNA in the centromere of chromosome XIV in Saccharomyces cerevisiae. Mol. Gen. Genet. 219, 153–160.

33. Dendooven, T., Zhang, Z., Yang, J., McLaughlin, S.H., Schwab, J., Scheres, S.H.W., Yatskevich, S., and Barford, D. (2023). Cryo-EM structure of the complete inner kinetochore of the budding yeast point centromere. Science advances 9. 10.1126/SCIADV.ADG7480.

34. Popchock, A.R., Hedouin, S., Mao, Y., Asbury, C.L., Stergachis, A.B., and Biggins, S. (2024). Stable centromere association of the yeast histone variant Cse4 requires its essential N-terminal domain. bioRxiv, 2024.07.24.604937. 10.1101/2024.07.24.604937.

35. Steinberg, K.M., Schneider, V.A., Graves-Lindsay, T.A., Fulton, R.S., Agarwala, R., Huddleston, J., Shiryev, S.A., Morgulis, A., Surti, U., Warren, W.C., et al. (2014). Single haplotype assembly of the human genome from a hydatidiform mole. Genome Res. 24, 2066–2076.

36. Logsdon, G.A., Rozanski, A.N., Ryabov, F., Potapova, T., Shepelev, V.A., Catacchio, C.R., Porubsky, D., Mao, Y., Yoo, D., Rautiainen, M., et al. (2024). The variation and evolution of complete human centromeres. Nature 629, 136–145.

37. Tachiwana, H., Kagawa, W., Shiga, T., Osakabe, A., Miya, Y., Saito, K., Hayashi-Takanaka, Y., Oda, T., Sato, M., Park, S.Y., et al. (2011). Crystal structure of the human centromeric nucleosome containing CENP-A. Nature 476, 232–235.

38. Hasson, D., Panchenko, T., Salimian, K.J., Salman, M.U., Sekulic, N., Alonso, A., Warburton, P.E., and Black, B.E. (2013). The octamer is the major form of CENP-A nucleosomes at human centromeres. Nat. Struct. Mol. Biol. 20, 687–695.

39. Canzio, D., Chang, E.Y., Shankar, S., Kuchenbecker, K.M., Simon, M.D., Madhani, H.D., Narlikar, G.J., and Al-Sady, B. (2011). Chromodomain-mediated oligomerization of HP1 suggests a nucleosome-bridging mechanism for heterochromatin assembly. Mol. Cell 41, 67–81.

40. Kiewisz, R., Fabig, G., Conway, W., Baum, D., Needleman, D., and Müller-Reichert, T. (2022). Three-dimensional structure of kinetochore-fibers in human mitotic spindles. Elife 11. 10.7554/ELIFE.75459.

41. O’Toole, E., Morphew, M., and Richard McIntosh, J. (2020). Electron tomography reveals aspects of spindle structure important for mechanical stability at metaphase. Mol. Biol. Cell 31, 184–195.

42. Dumont, M., Gamba, R., Gestraud, P., Klaasen, S., Worrall, J.T., De Vries, S.G., Boudreau, V., Salinas-Luypaert, C., Maddox, P.S., Lens, S.M.A., et al. (2020). Human chromosome-specific aneuploidy is influenced by DNA-dependent centromeric features. EMBO J. 39. 10.15252/EMBJ.2019102924.

43. Vafa, O., and Sullivan, K.F. (1997). Chromatin containing CENP-A and alpha-satellite DNA is a major component of the inner kinetochore plate. Curr. Biol. 7, 897–900.

44. Kornberg, R.D. (1974). Chromatin structure: a repeating unit of histones and DNA. Science 184, 868–871.

45. Valouev, A., Johnson, S.M., Boyd, S.D., Smith, C.L., Fire, A.Z., and Sidow, A. (2011). Determinants of nucleosome organization in primary human cells. Nature 474, 516–522.

46. Ikeno, M., Masumoto, H., and Okazaki, T. (1994). Distribution of CENP-B boxes reflected in CREST centromere antigenic sites on long-range α-satellite DNA arrays of human chromosome 21. Hum. Mol. Genet. 3, 1245–1257.

47. Yoda, K., Ando, S., Okuda, A., Kikuchi, A., and Okazaki, T. (1998). In vitro assembly of the CENP-B/alpha-satellite DNA/core histone complex: CENP-B causes nucleosome positioning. Genes Cells 3, 533–548.

48. Tanaka, Y., Tachiwana, H., Yoda, K., Masumoto, H., Okazaki, T., Kurumizaka, H., and Yokoyama, S. (2005). Human centromere protein B induces translational positioning of nucleosomes on α-satellite sequences. J. Biol. Chem. 280, 41609–41618.

49. Tanaka, Y., Kurumizaka, H., and Yokoyama, S. (2005). CpG methylation of the CENP-B box reduces human CENP-B binding. FEBS J. 272, 282–289.

50. Okada, T., Ohzeki, J.I., Nakano, M., Yoda, K., Brinkley, W.R., Larionov, V., and Masumoto, H. (2007). CENP-B Controls Centromere Formation Depending on the Chromatin Context. Cell 131, 1287–1300.

51. Tanaka, Y., Nureki, O., Kurumizaka, H., Fukai, S., Kawaguchi, S., Ikuta, M., Iwahara, J., Okazaki, T., and Yokoyama, S. (2001). Crystal structure of the CENP-B protein-DNA complex: the DNA-binding domains of CENP-B induce kinks in the CENP-B box DNA. EMBO J. 20, 6612–6618.

52. Earnshaw, W.C., Sullivan, K.F., Machlin, P.S., Cooke, C.A., Kaiser, D.A., Pollard, T.D., Rothfield, N.F., and Cleveland, D.W. (1987). Molecular cloning of cDNA for CENP-B, the major human centromere autoantigen. J. Cell Biol. 104, 817–829.

53. Capozzi, O., Carbone, L., Stanyon, R.R., Marra, A., Yang, F., Whelan, C.W., De Jong, P.J., Rocchi, M., and Archidiacono, N. (2012). A comprehensive molecular cytogenetic analysis of chromosome rearrangements in gibbons. Genome Res. 22, 2520–2528.

54. Hartley, G.A., Okhovat, M., O’Neill, R.J., and Carbone, L. (2021). Comparative Analyses of Gibbon Centromeres Reveal Dynamic Genus-Specific Shifts in Repeat Composition. Mol. Biol. Evol. 38, 3972–3992.

55. Walfridsson, J., Bjerling, P., Thalen, M., Yoo, E.J., Park, S.D., and Ekwall, K. (2005). The CHD remodeling factor Hrp1 stimulates CENP-A loading to centromeres. Nucleic Acids Res. 33, 2868–2879.

56. Tawaramoto, M.S., Park, S.Y., Tanaka, Y., Nureki, O., Kurumizaka, H., and Yokoyama, S. (2003). Crystal structure of the human centromere protein B (CENP-B) dimerization domain at 1.65-A resolution. J. Biol. Chem. 278, 51454–51461.

57. Chardon, F., Japaridze, A., Witt, H., Velikovsky, L., Chakraborty, C., Wilhelm, T., Dumont, M., Yang, W., Kikuti, C., Gangnard, S., et al. (2022). CENP-B-mediated DNA loops regulate activity and stability of human centromeres. Mol. Cell 82, 1751–1767.e8.

58. Henikoff, S., Ahmad, K., and Malik, H.S. (2001). The centromere paradox: stable inheritance with rapidly evolving DNA. Science 293, 1098–1102.

59. Logsdon, G.A., and Eichler, E.E. (2022). The Dynamic Structure and Rapid Evolution of Human Centromeric Satellite DNA. Genes 14, 92.

60. Grant, C.E., Bailey, T.L., and Noble, W.S. (2011). FIMO: Scanning for occurrences of a given motif. Bioinformatics 27, 1017–1018.

61. Kolmogorov, M., Yuan, J., Lin, Y., and Pevzner, P.A. (2019). Assembly of long, error-prone reads using repeat graphs. Nat. Biotechnol. 37, 540–546.

62. Carbone, L., Alan Harris, R., Gnerre, S., Veeramah, K.R., Lorente-Galdos, B., Huddleston, J., Meyer, T.J., Herrero, J., Roos, C., Aken, B., et al. (2014). Gibbon genome and the fast karyotype evolution of small apes. Nature 513, 195–201.

63. Okhovat, M., Nevonen, K.A., Davis, B.A., Michener, P., Ward, S., Milhaven, M., Harshman, L., Sohota, A., Fernandes, J.D., Salama, S.R., et al. (2020). Co-option of the lineage-specific LAVA retrotransposon in the gibbon genome. Proc. Natl. Acad. Sci. U. S. A. 117, 19328–19338.

64. Durand, N.C., Shamim, M.S., Machol, I., Rao, S.S.P., Huntley, M.H., Lander, E.S., and Aiden, E.L. (2016). Juicer Provides a One-Click System for Analyzing Loop-Resolution Hi-C Experiments. Cell systems 3, 95–98.

65. Dudchenko, O., Batra, S.S., Omer, A.D., Nyquist, S.K., Hoeger, M., Durand, N.C., Shamim, M.S., Machol, I., Lander, E.S., Aiden, A.P., et al. (2017). De novo assembly of the Aedes aegypti genome using Hi-C yields chromosome-length scaffolds. Science 356, 92–95.

